# Prion shedding is reduced by chronic wasting disease vaccination

**DOI:** 10.64898/2025.12.16.694724

**Authors:** Hanaa Ahmed-Hassan, Dalia Abdelaziz, Yo-Ching Cheng, Kevin Low, Shirley Phan, Byron Kruger, Chimoné S. Dalton, Lech Kaczmarczyk, Walker S. Jackson, Sabine Gilch, Hermann M. Schätzl

## Abstract

Chronic wasting disease (CWD) is a strictly fatal and highly contagious prion disease of wild and farmed cervids currently expanding in North America. Prion diseases are caused by conversion of the cellular prion protein to its pathological isoform PrP^Sc^. Vaccination is considered a promising strategy to contain CWD, even though prion diseases do not show classical immune responses. For CWD containment, it is important that vaccines reduce shedding of prions in excreta, a major contributor to transmission. Here, we tested the effect of vaccines on prion shedding in feces and urine by vaccinating and prion infecting knock-in mice that recapitulate CWD pathogenesis as found in cervids. Vaccination reduced or even prevented CWD shedding in feces and urine collected between 30-90% of incubation time to disease. This is the first report showing that prion shedding can be blocked in a prion disease. For CWD specifically it may reduce the environmental prion burden and break the disease transmission cycle.

## Introduction

Prion diseases or transmissible spongiform encephalopathies (TSE’s) are fatal infectious neurodegenerative disorders of the central nervous system (CNS). These diseases are caused by the conversion of the normal cellular prion protein (PrP^C^) into the misfolded and pathological scrapie isoform termed PrP^Sc^ ^1^. Accumulation of PrP^Sc^ leads to spongiform degeneration and neuronal loss in the CNS ^2^. Chronic Wasting Disease (CWD) is considered the most contagious prion disease in animals, affecting both wild and captive cervid populations such as elk, deer, reindeer and moose ^3–5^. CWD is expanding rapidly in North America, and by December 2025 it has been detected in 36 U.S. states and 5 Canadian provinces ^3^. CWD has also been found in South Korea and since 2016 in Norway, Finland, Sweden ^3,6^.

CWD is the only known prion disease that affects wildlife, which is a significant challenge for controlling the disease. Prion infectivity is present in the CNS as well as many peripheral tissues, body fluids and secretions of CWD infected animals ^7, 8^. Transmission of CWD occurs directly or indirectly between animals, due to shedding of CWD prions through feces, saliva and urine, which contaminates the environment. Importantly, shedding starts early in pre-clinical disease stages ^9–12^. Environmental contamination of CWD persists for years and serves as a reservoir for infecting the same species, or it may facilitate transmission to other susceptible species. CWD is economically relevant and can directly impact hunting and tourism industries, and potentially food safety and food security of people who rely on subsistence hunting and the consumption of venison. The uncertain zoonotic potential of CWD remains a significant risk for humans ^13^. CWD can be transmitted experimentally to non-cervid hosts including voles, hamsters, ferrets, sheep, cats, mink, pigs and cattle ^14–18^. Several studies that examined the zoonotic potential of CWD concluded that the risk is very low ^19–24^. However, a recent study showed that CWD can be transmitted to mice expressing the human prion protein, with CWD infectivity found in brain and feces ^20^. If this disease manifestation holds true for humans, it would make feces a source of infection between humans. In addition, CWD prions have been detected in skeletal muscle ^25^ and antler velvet of infected cervids, which raises the possibility of zoonotic transmission through consumption of contaminated venison and/or utilization of infected cervid products in Asian medicines ^26^.

To date, there is no effective treatment or vaccination available against CWD. Prion disease does not evoke a detectable immune reaction in the host, as the endogenous host PrP^C^, a self-protein, is converted into PrP^Sc^ when prions start to replicate. The main obstacle to having a vaccine against any prion disease is therefore overcoming the self-tolerance without inducing unwanted side effects ^5^. Conceptually there are two main targets for active vaccination against prion diseases: targeting PrP^C^ or PrP^Sc 5, 27–30^. The approach pioneered by our group targets PrP^C^ (termed ‘PrP^C^ vaccination’) and mostly induces self-antibodies that bind to surface-located PrP^C^ and interfere with cellular prion replication ^31–33^. Ideally, an effective vaccine for CWD should decrease the number of infected cervids over time, reduce the shedding of already infected cervids, and reduce the contamination of the environment.

Our previous studies have shown that vaccination targeting PrP^C^ overcame self-tolerance in cervids and transgenic mice, produced detectable humoral and cellular immune responses, and increased the survival time in mouse models of CWD without inducing unwanted side effects ^5, 34–36^. No study has yet tested the efficacy of active vaccination on CWD shedding. In the current study, we show the effect of vaccination on CWD shedding in feces and urine, using cervidized knock-in (KI) mice that recapitulate CWD pathogenesis as found in the cervid host ^37^. We used recombinant dimeric deer prion protein (Ddi) and monomeric mouse prion protein (Mmo), vaccines we have shown to produce humoral immune responses against PrP^C 5,31,33^, and able to protect vaccinated cervidized mice against prion challenge ^34–36^. To investigate the effect of Ddi and Mmo vaccination on CWD shedding, feces and urine were collected every 50 days post infection (dpi). Fecal samples contain inhibitors that make CWD detection more difficult by in vitro amplification assays such as protein misfolding cyclic amplification (PMCA) and real-time quaking-induced conversion (RT-QuIC) ^38–41^. To improve the sensitivity of those assays, iron oxide magnetic extraction (IOME) beads have been used ^42, 43^. In the current study, we combine IOME with PMCA followed by RT-QuIC to increase sensitivity and specificity of the assay. The combination of the three techniques is referred here as ‘IPR technique’. Our study demonstrates that vaccination targeting PrP^C^ leads to a decrease in CWD shedding in feces and urine at preclinical stages in a CWD mouse model. This vaccination strategy therefore has two additive effects: it improves individual survival which will translate into population effect, and it reduces prion shedding translating into reduction of CWD prions in the environment.

## Results

### Vaccination targeting PrP^C^ delays CWD neuro-invasion in knock-in mice inoculated intraperitoneally with CWD

In this study, we tested the effect of PrP^C^ vaccination on CWD prion shedding in feces and urine using a cervidized KI mouse model. We used gene-targeted KI mice where mouse PrP was replaced with cervid PrP, which recapitulate the pathogenesis of CWD, including shedding of CWD prions into feces and urine ^37^. For vaccination, we used three groups of KI mice (n=8/group): two vaccine groups receiving either recombinant dimeric deer (Ddi) or monomeric mouse (Mmo) PrP with CpG as adjuvant, or CpG adjuvant alone as control group. All mice received one priming dose (100 µg of the protein plus 5 µM of CpG) and four booster doses subcutaneously at three weeks intervals (s.c.) (50 µg of the protein plus 5 µM of CpG). The control group received 5 µM CpG in PBS only. CWD prion infection was performed intraperitoneally (i.p.) using brain homogenate of KI-mouse adapted reindeer CWD **(Fig. 1a)**. This mouse model was expected to develop clinical signs of prion infection between 450 and 500 dpi ^37^. We harvested various mice before reaching the clinical endpoint and used such pre-clinical mice to assess differences in PrP^Sc^ levels in brain and spinal cord between immunized and control mice by immunoblot analysis. Overall, PrP^Sc^ levels in brain **(Fig. 1c, d)** and spinal cord **(Fig. 1e, f)** of mice with comparable incubation time in the Ddi or Mmo vaccinated groups were reduced compared to control animals. As expected, PrP^Sc^ levels in spinal cord were higher than in brain, reflecting the peripheral mode of infection and the process of neuro-invasion **(Fig. 1g-i)**. Only five mice reached terminal prion disease and presented clinical signs such as belly movement, extension, and clasping of hind limbs. One CpG (#6) mouse showed clinical signs of terminal prion disease at 462 dpi, while two mice from the Mmo (#2 and 4) vaccinated group developed terminal prion disease at 494 and 495 dpi, and two mice in the Ddi (#2 and 3) vaccinated group at 488 and 495 dpi, respectively. PrP^C^ vaccination increased survival time by around 30 days compared to the control group, although a statistical analysis is not possible.

**Fig. 1.**
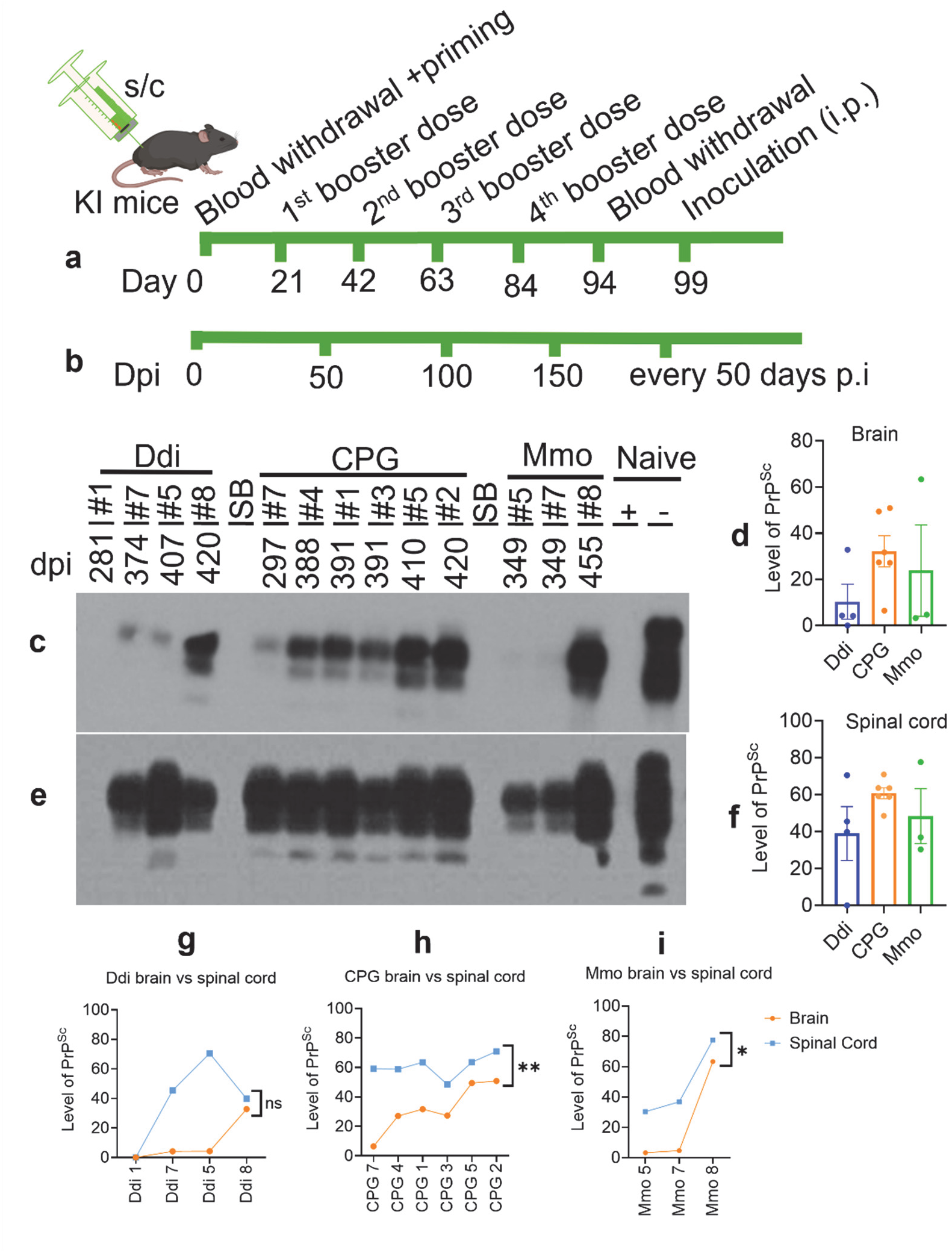
Immunoblot showing PrP^Sc^ levels in brain and spinal cord of cervid KI mice. **a,** Schematic diagram showing the study design for vaccination and **b,** the time points of feces and urine collection. Ddi vaccinated group, CPG control group and Mmo vaccinated group inoculated i.p. with 1% brain homogenate (BH) of mouse-adapted reindeer CWD. Samples were digested with 50 µg/ml PK and loaded onto SDS-PAGE as labelled at the top of the blot **c,** brain and **e,** spinal cord. Membranes were probed with the 4H11 antibody (1:500). Naive: brain of non-inoculated KI mice. The PrP^res^ level in **c,** brain, **e,** Spinal cord, **g,** brain vs spinal cord in Ddi group, **h,** brain vs spinal cord in CPG group, and **I,** brain vs spinal cord in Mmo group. The delay in the neuro-invasion in the vaccinated groups represented in graph **c** and **e**. Graphs generated by GraphPad Prism (version 10). Statistical analysis done using paired-t test, ns=not significance, * P-value= 0.0440 and ** P-value= 0.0038.

These data show that vaccination delayed the process of neuro-invasion and extended the time to clinical disease in both vaccinated groups compared to the control group.

### PrP^C^ vaccination induces a humoral immune response in KI mice

For testing specific humoral immune responses, we analyzed post-immune sera from all mice in ELISA, using plates coated with Ddi as antigen **(Fig. 2a, b)** as well as plates coated with Mmo **(Fig. 2c, d)**. We found that all mice vaccinated with Ddi immunogen showed reactivity against Ddi antigen (**Fig. 2a, b**), but not against Mmo (**Fig. 2c, d**) at a 1:100 dilution. Mice vaccinated with Mmo immunogen reacted well in both situations. The endpoint dilution of sera showed that most sera were still reactive at a 1:30,000 dilution **(Fig. 2e)**. These data show that both immunogens break the self-tolerance to PrP and produce high antibody titers. To gain a deeper understanding of the specificity of the immune responses generated by the vaccine candidates, we performed a linear epitope mapping analysis, covering full-length mature cervid PrP. The post-immune sera used in this analysis were average from 4 mice in the Ddi-vaccinated group (mice # 1, 3, 4 and 6) and 4 mice from the Mmo group (1, 2, 3 and 6), so it reflects average reactivity in each group. Of note, there was no reactivity to the polyhistidine tag (polyhis epitope) and the linker sequence **(Extended Data Fig.1)**. Interestingly, both immunogens resulted in a similar reactivity to the linear epitopes, with exception of epitope #11 (sequence:NTFVHDCVNITVKQHTVTTTT) (**Extended Data table 1)**, to which Ddi-vaccinated sera reacted but not Mmo sera **(Extended Data Fig.1)**.

**Fig. 2.**
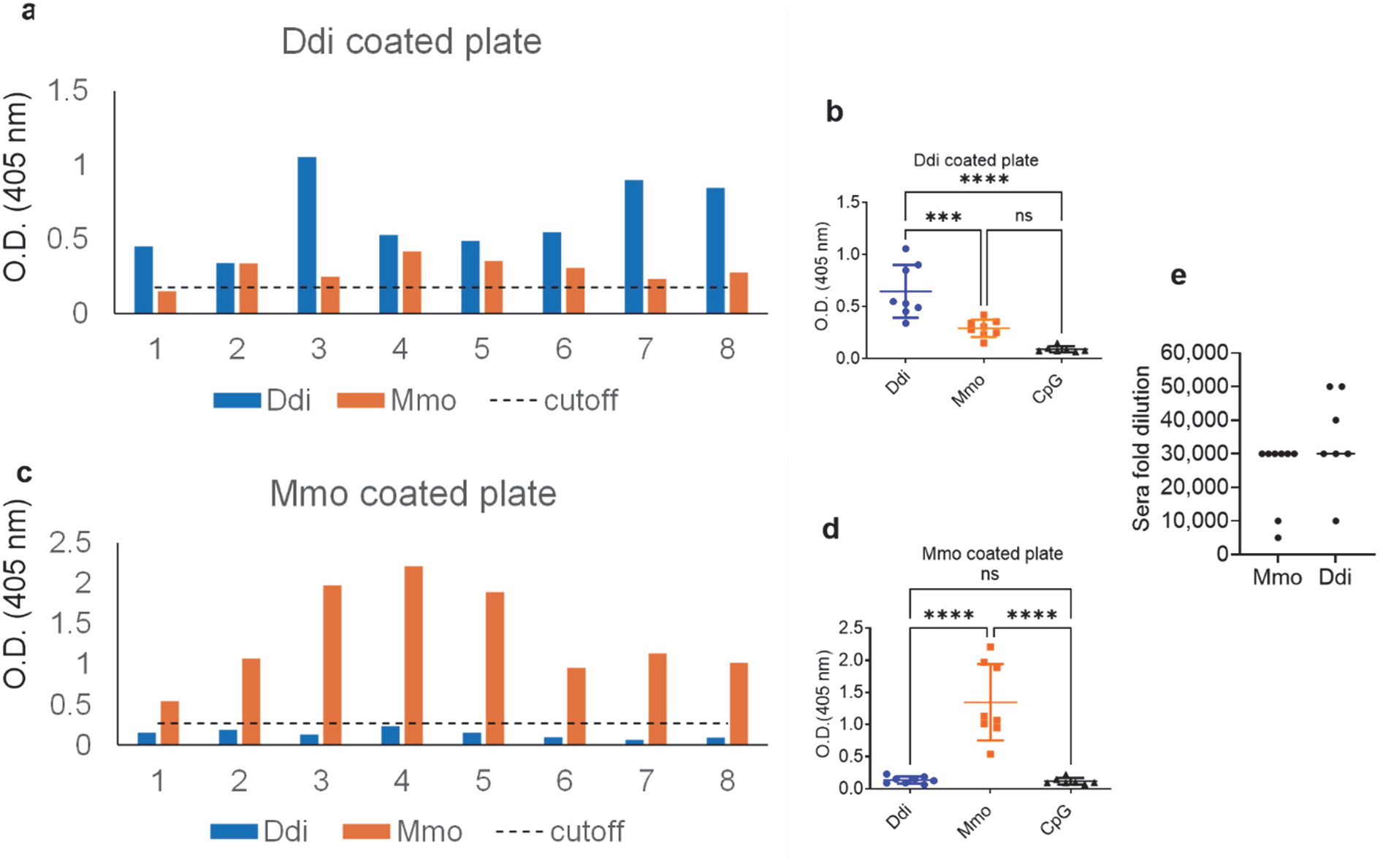
Humoral immune response of Ddi & Mmo vaccinated KI mice to immunogens. ELISA plates were coated with **a-b,** Ddi (immunogen) or **c-d,** Mmo (immunogen). Sera was isolated after the 4^th^ dose of vaccination. All sera were diluted at 1:100 and CpG mice sera were used as a negative control. One-way ANOVA followed by Tukey’s test. Statistical significance = **** P-value <0.0001, *** P-value=0.0007. **e,** End-point ELISA-purposed antibody titers are shown for the two vaccinated groups. Each mouse antibody titer was determined by endpoint dilution and represented by a data point. The y-axis represents the serum fold dilution, and the X-axis represents the treated groups. The dashed horizontal line represents the cut-off, which is the average OD of CpG sera +3*SD.

### Vaccination decreases CWD prion shedding in feces of KI mice

To test the effect of vaccination on shedding of CWD prions in feces, individual and pooled fecal samples were collected starting at 200 dpi. Fecal samples were subjected to homogenization and iron oxide magnetic extraction (IOME), followed by protein misfolding cyclic amplification (PMCA) and real-time quaking-induced conversion (RT-QuIC), summarized here as IPR readout. Of note, this is the first description of IPR to detect CWD in mouse feces. First, we started screening pooled fecal samples taken at 350 dpi. IPR, with RT-QuIC as final readout, showed that the seeding activity in the Ddi group was lower than that of the CpG and Mmo groups. Pooled fecal samples from one cage with Ddi-vaccinated mice showed no reactivity **(Fig. 3a**), whereas pooled samples from the second cage had reactivity, but lower than that of CpG control mice or Mmo-vaccinated mice (**Fig. 3b**). To get a better resolution of this finding in time and at the individual mouse level, we next studied fecal samples of individual mice. For this, we analyzed samples taken at 200, 250, 300 and 350 dpi, representing 45%, 55%, 65% and 75% of the incubation time. We observed that CWD shedding into feces can be detected as early as 200 dpi in KI mice. Interestingly, we found that only 50% of fecal samples from Ddi-vaccinated mice at 200 dpi tested positive, whereas samples from Mmo-vaccinated mice were positive at 87.5%, and samples of CPG control mice at 100% **(Fig. 4a-d**, **Table 1)**. In addition, samples from Ddi- and Mmo-vaccinated mice still testing positive showed overall a lower seeding activity, as determined by a higher time to threshold **(Fig. 4e)**, lower maximum of range **(Fig. 4f)**, and lower area under the curve **(Fig. 4g)** analysis. These data show that vaccination with Ddi and Mmo immunogens reduced shedding of CWD prions into feces at 200 dpi. Similar results were obtained at 250, 300 and 350 dpi. At 250 dpi, four out of seven mice (57%) tested positive for shedding in the control group versus two out of seven (28%) in the Ddi-immunized group **(Extended Data Fig. 2, Table 1)**. At 300 dpi, a similar trend was found, with two out of five mice (40%) testing positive in Ddi-vaccinated mice, but 100% positivity in the CpG control group **(Extended Data Fig. 3, Table 1)**. Fecal samples of Ddi-vaccinated mice that were still positive had lower quantities of CWD prions in the feces taken at 200, 250 and 300 dpi **(Fig 4, Extended Data Fig. 2 and 3 e-g**), with a similar trend for fecal samples from Mmo-vaccinated mice. Interestingly, at 350 dpi, the positivity rate was lowest in the Mmo-vaccinated group (20%), compared to Ddi-vaccinated mice (66%) CpG-only treated control mice (71%) (**Extended Data Fig. 4 a-d**, **Table 1**). As before, samples still being reactive showed less signal intensity when analyzing time to threshold, area under the curve or maximum range compared to CpG-only treated mice, now more pronounced for Mmo-vaccinted mice **(Extended Data Fig. 4 e-g)**. When correlating shedding results with antibody titers, the Ddi sera of mice #6, 7 and 8 showed the highest antibody titers, with 1:40,000 and 1:50,000, respectively. Ddi mice #7 and 8 showed shedding activity only at one of the four time points analyzed **(Extended Data Table 3)**. In addition, Ddi mice #7 and 8 showed low PrP^Sc^ levels in spinal cord. On the contrary side, Mmo mouse #8 has a low antibody titer (1:5,000) and high PrP^Sc^ levels in brain and spinal cord, when compared to Mmo mice #5 and 7 with higher titers (1:30,000) (**Fig 1 c, e and Extended Data Table 3**).

**Fig. 3.**
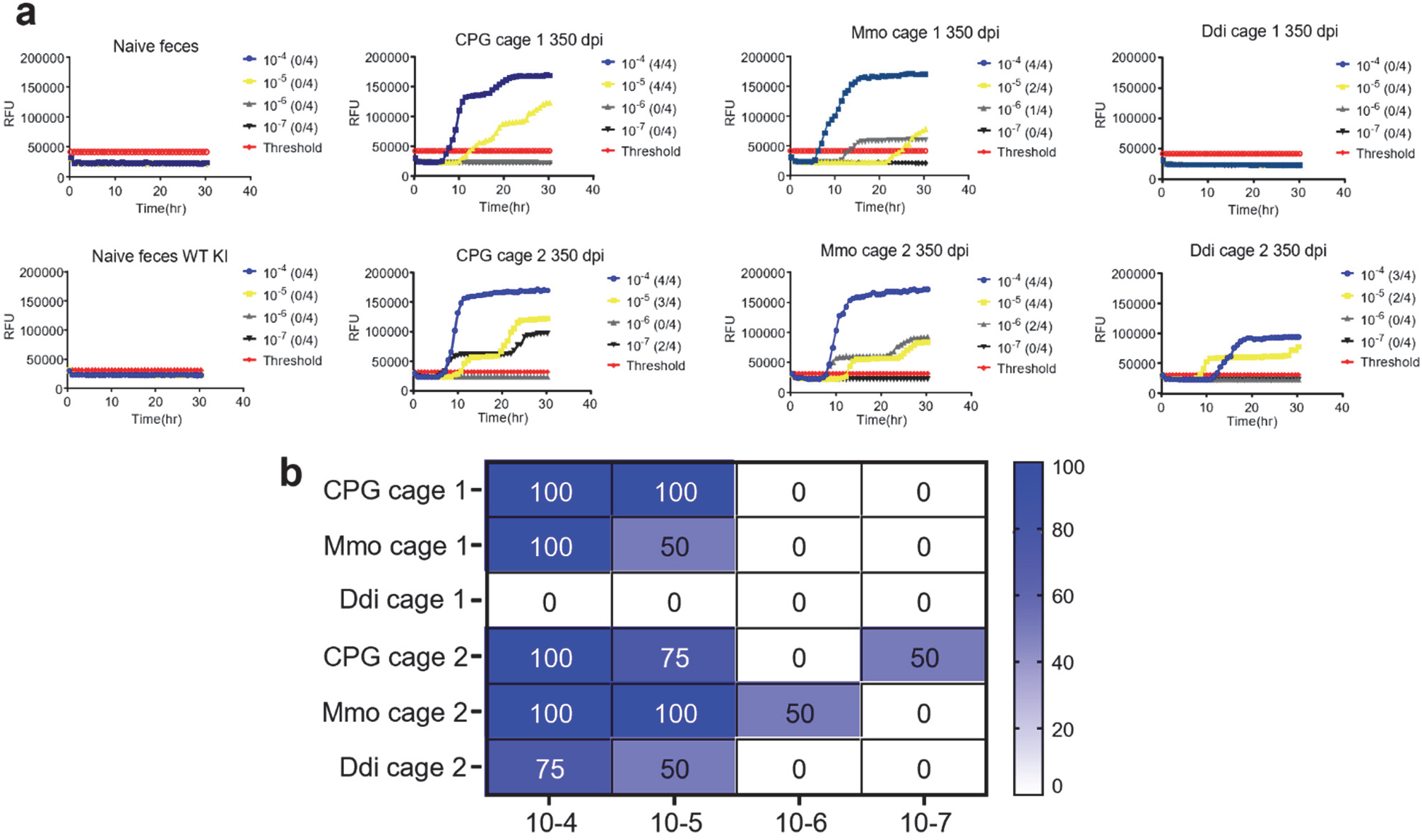
RT-QuIC data for the PMCA products showing the difference in the CWD seeding activity between KI vaccinated and control groups at 350 dpi. 10% pooled fecal homogenate from all groups (two cages per group) extracted using IOME followed by three rounds of PMCA reactions using KI WT138 PrP^C^ substrate seeded with 1:10 dilution of the fecal homogenate, in duplicates for all vaccinated and control group. Positive control for the PMCA was naïve feces spiked with mouse-adapted CWD reindeer; negative control was naïve feces. Samples and controls were subjected to 3 rounds of PMCA. The PMCA products analyzed using RT-QuIC assay at 10^-4^ to 10^-7^ dilution. **a,** Representative RT-QuIC graphs showing the seeding activity in feces from KI WT138 mice vaccinated subcutaneous with Ddi or Mmo or injected with CPG as control followed by intraperitoneally inoculation with mouse-adapted CWD Reindeer respectively. Samples were considered positive when 2 out of 4 wells crossed the threshold, which defined as the average relative fluorescent unit (RFU) of the negative control group plus five times its standard deviation. The y-axis represents the RFU, and the x-axis represents the time in hours(hr). **b,** Summary of the RT-QuIC analysis of prion seeding in the 350dpi pooled feces of KI mice. The heat map indicates the percentage of positive RT-QuIC replicates out of the total of four replicates analyzed. The scale ranges from 0 (all replicates were negative) to 100 (all replicates were positive). Graphs were generated using GraphPad Prism (version 10).

**Fig. 4.**
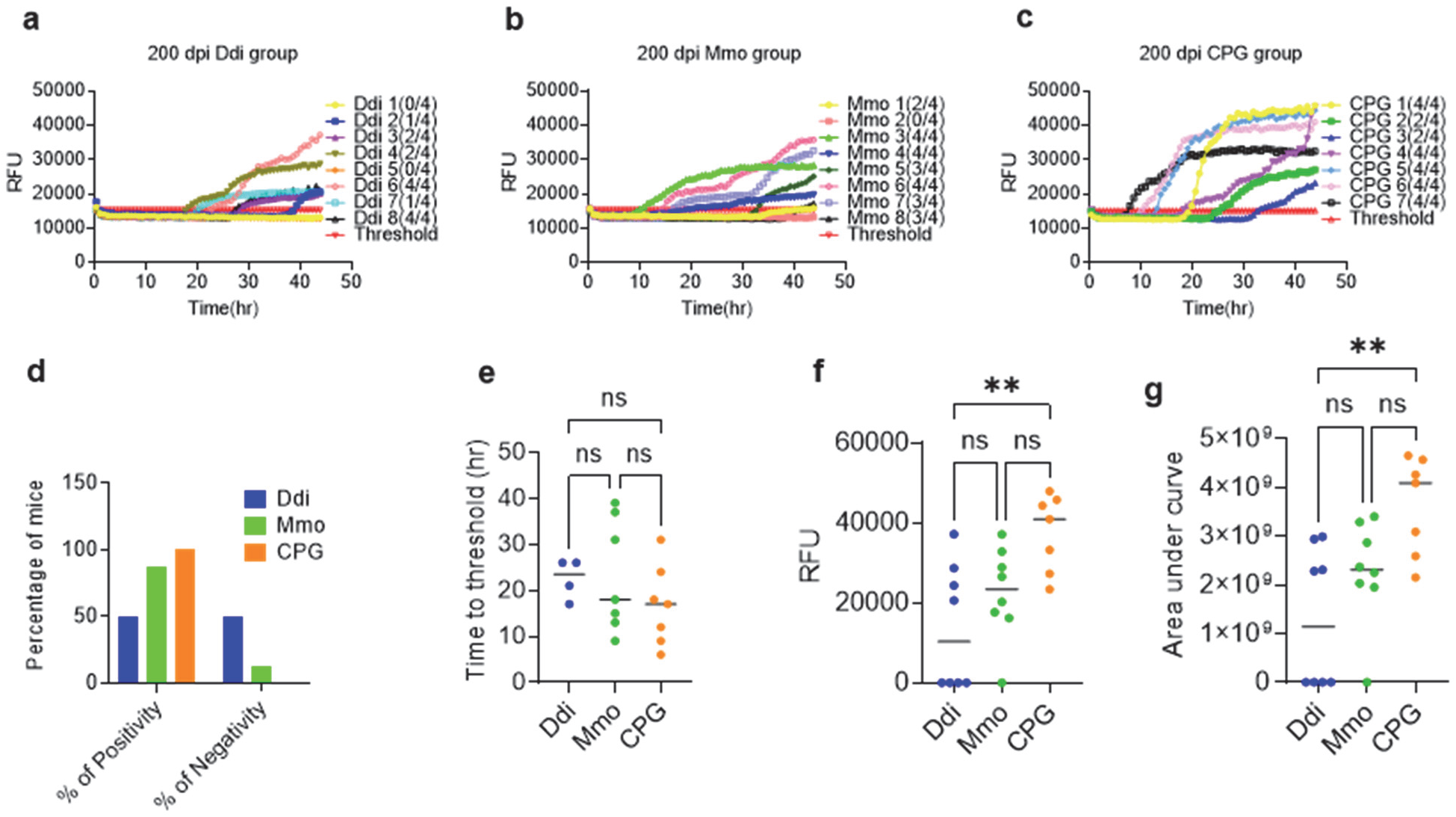
Seeding activity in individual fecal samples of KI vaccinated and control group at 200 dpi. Three rounds of PMCA reactions were done using KI substrate seeded with individual 10^-1^ dilution of 200 dpi 10% fecal homogenate KI WT 138 after IOME, in duplicates for all vaccinated and control groups. Positive control for the PMCA was naïve feces spiked 1:100 with mouse-adapted CWD Reindeer and negative control was naïve feces subjected to three cycles of PMCA reaction. The PMCA products analyzed using RT-QuIC assay at 10^-1^ dilution. **a-c,** Representative RT-QuIC graphs showing the seeding activity in feces from KI WT138 mice vaccinated with Ddi or Mmo and CPG control group. Samples were considered positive when 2 out of 4 wells crossed the threshold, which defined as the average RFU of the negative control group plus five times its standard deviation. The y-axis represents the RFU, and the x-axis represents the time in hours(hr). **d,** Chi square test, **e,** Time to threshold, **f,** Maximum of range and **g,** Area under curve. Graphs were generated using GraphPad Prism (version 10). Statistical analysis done using Chi-square test and **** p-value< 0.0001- or One-way ANOVA followed by a Tukey’s multiple comparison. For Maximum of range ** p-value= 0.0046, For area under curve ** p-value= 0.0034 and ns: not significant.

**Table 1:** Difference in the CWD shedding in feces between KI vaccinated and control groups at different time points.

|  | **% of positivity** |  |  |  |
| --- | --- | --- | --- | --- |
| **Dpi** | **200 dpi** | **250 dpi** | **300 dpi** | **350 dpi** |
| **Ddi** | (4/8) 50% | (2/7) 28.6% | (2/5) 40% | (4/6) 66.6% |
| **Mmo** | (7/8) 87.5% | (4/7) 57.1% | (5/6) 83.3% | (1/5) 20% |
| **CPG** | (7/7) 100% | (4/7) 57.1% | (5/5) 100% | (5/7) 71.4% |

Taken together, fecal samples from individual Ddi- and Mmo-vaccinated mice showed lower positivity rates than samples from CpG control mice, at 200, 250, 300 and 350 dpi (**Table 1**). In addition, CpG samples consistently showed the highest prion seeding activities compared to the vaccinated samples. This shows that vaccination consistently reduced shedding of CWD prions into feces, resulting in less positive fecal samples and in lower CWD prion amounts in still positive samples.

### Vaccination with Ddi immunogen blocks shedding of CWD prions in urine of KI mice

Having seen a consistent reduction of CWD prion shedding in the feces of vaccinated mice, we next wanted to see whether shedding is also reduced in the urine of vaccinated mice at various preclinical stages. To complement our shedding analysis in feces, which analyzed samples taken between 200 and 350 dpi only, we chose for the urine analysis also earlier (150 dpi, corresponding to 30% of incubation time) and later samples (450 dpi, corresponding to 90% of incubation time). Since it was technically not possible to analyze urine samples from individual mice due to the small volumes available, we decided to pool urine samples. Urine samples were analyzed using IOME extraction followed by three rounds of PMCA, or the IPR technique as described above. Remarkably, Ddi and Mmo-immunized mice did not shed CWD prions in the urine at 150, 250 and 450 dpi, when coupling IOME extraction and PMCA with immunoblot read-out **(Fig. 5a, b)**. CWD seeding activity was also not detected in urine samples of Ddi and Mmo-vaccinated mice at 150 and 250 dpi when performing the more sensitive IPR analysis as revealed in the RT-QuIC heat map (**Fig. 5c, d**). At both time points, there was no detectable reactivity in urine samples of vaccinated mice, with a 4-log higher titer in urine samples of CpG control mice, where positivity was found at 10^-1^ to 10^-4^ dilutions of RT-QuIC reactions. These results indicate a strong presence of prions in urine of control mice, which was drastically reduced in vaccinated mice (**Fig. 5 c,d and Extended Data fig. 5**). At 450 dpi, we had samples from 6 pre-clinical mice in total only **(Extended Data table 2)**. Samples from the 3 remaining Ddi-vaccinated mice (#2-4) were negative in all RT-QuIC dilutions, indicating no detectable shedding in urine. Samples from Mmo-vaccinated mice (n=2, #2 and 4) were now positive up to the 10^-4^ dilution. Samples from the remaining CpG control mouse (#6) tested positive up to the 10^-5^ dilution (**Fig. 5e and Extended Data Fig 6**), indicating a 5-log reduction of shedding in Ddi-vaccinated mice.

**Fig. 5.**
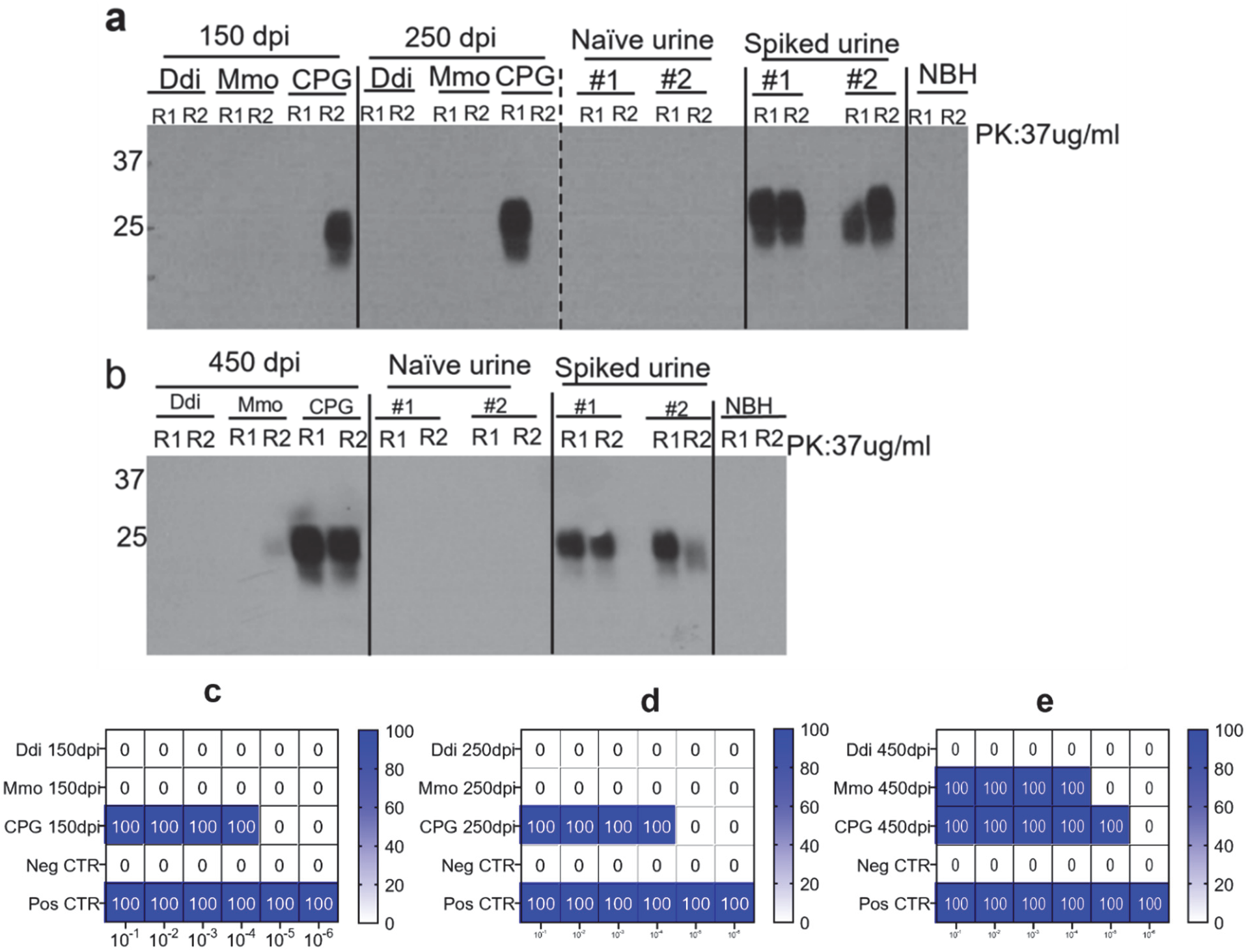
Representative Western blots showing PrP^Sc^ levels in the third round of PMCA reactions using wild type cervid PrP^C^ substrate seeded with 10^-1^ dilution of urine of Ddi, Mmo vaccinated or CPG control KI mice in duplicates at different time points. **a**, 150 and 250 dpi and **b,** 450 dpi samples were subjected to IOME followed by three rounds of PMCA. Negative control for the PMCA is naïve urine and positive control for the PMCA was naïve urine spiked (1:100) with brain homogenate of mouse-adapted CWD Reindeer subjected to PMCA reaction. PMCA products were digested for 1 hour with 37 µg/ml PK at 37°C and were probed with the 4H11 antibody (1:500). NBH: normal brain homogenate, which is the substrate, used in the PMCA as another control. **c-e,** Summary of the RT-QuIC analysis of prion seeding in the pooled urine of KI mice at **c,** 150 dpi, **d,** 250 dpi, and **e,** 450 dpi. The heat map indicates the percentage of positive RT-QuIC replicates out of the total of four replicates analyzed. The scale ranges from 0 (all replicates were negative) to 100 (all replicates were positive). Graphs were generated using GraphPad Prism (version 10).

Taken together these results indicate that vaccination with the Ddi immunogen can prevent shedding of CWD prions in urine in the KI mouse model of CWD infection.

## Discussion

This is the first study which shows that vaccination reduces the shedding of CWD prions into urine and feces. This is a very important finding for the future containment of CWD, both for farmed and wild-living animals. A vaccination that extends the lifetime of CWD-infected cervids without reducing the shedding of prions would be counter-productive, increasing the environmental load with prions and likely not breaking the disease transmission cycle.

CWD is a highly contagious disease that can be transmitted directly or indirectly between cervids. CWD-infected cervids contaminate the environment through continuous shedding of CWD prions in secretions such as feces, saliva, and urine, both in clinical and early pre-clinical stages ^43–45^. Prions in the environment are very stable and can remain infectious over years. Experimental studies showed that prion seeding activity of cervid feces can remain detectable despite environmental factors such as desiccation and freeze-thaw cycles ^41^. CWD prions effectively attach to certain soil types and are absorbed by plants, potentially contaminating the environment for decades ^46–48^, making disease containment and disease control very challenging. Horizontal transmission of CWD prions between cervids occurs through the alimentary tract that is the major route of entry for CWD prions ^49^. From there, the process of neuro-invasion is slow and provides a window of opportunity for vaccination strategies, as antibodies do not have to cross the blood–brain barrier to be effective ^29,30,50^. When the process of neuro-invasion is established, CWD prions egress from the central nervous system in a process not well understood and finally are found in peripheral tissues like muscles and secretions like urine, feces and saliva. Again, antibodies might have the opportunity to interfere in this process, acting outside the central nervous system ^5,36,51^.

Importantly, prion infections do not evoke detectable immune responses. In contrast to traditional infectious agents that are ‘foreign’ to the immune system, prions use endogenous PrP^C^ for their replication, which is a self-protein to the immune system. This also makes newly produced PrP^Sc^ a self-protein. The challenge for vaccine strategies is to effectively overcome this self-tolerance but doing so without inducing unwanted side effects. We and others have shown experimentally that this is possible ^27–35^. Conceptually, there are two targets for active vaccination against prion diseases: PrP^C^, the substrate needed for prion conversion, or PrP^Sc^ that acts as template ^5, 36^ In both situations, binding of anti-PrP antibodies might be sufficient to interfere by sterical hindrance with prion propagation, a process that involves a fitted protein-protein interaction. The PrP^Sc^-targeted approach uses disease-specific epitopes (DSE), which are not present in PrP^C^ and only are accessible in forming or already formed PrP^Sc^. Several DSEs have been described: YYR, YML, and RL. The rigid loop region (RL) elicits systemic and mucosal immune responses when administered orally to white-tailed deer ^52^. However, experimental trials in large animals using a YYR-based vaccine are inconsistent. In a sheep model challenged orally, the vaccine delayed the onset of disease ^53^, but the disease was accelerated in elk when they were exposed to a CWD-contaminated environment for a long period ^54^. Another group used a VPrP^Sc^ vaccine, recombinant antigens that mimic a 4-rung beta solenoid fold of PrP^Sc^, which delayed the onset of symptoms in a transgenic mouse model of a genetic human prion disease ^55^. However, no studies have been reported with this vaccine in CWD models.

The PrP^C^-targeting approach developed by us results in humoral and cellular immune responses against the immunogen without unwanted side effects ^5,31,33^. Induced self-antibodies target surface-located PrP^C^, which leads to depletion of PrP^C^ and in blocking its role as a substrate for prion conversion ^56,57^. This was shown by us in mouse and cervid models ^31,33–35^. Using transgenic mice (Tg) expressing cervid PrP we have shown that vaccination prolongs the survival time by up to 70% upon intraperitoneal CWD challenge ^35^. Unfortunately, such trangenic mice do not completely recapitulate the pathogenesis as found in the real cervid host. For instance, they do not shed prions into urine and feces ^5,58^. To overcome this problem, we used gene-targeted KI mice in this study, for which it was shown that they closely recapitulate CWD pathogenesis, including shedding ^59^. This allows us to investigate whether vaccination affects shedding of CWD prions, which was not feasible in transgenic mouse models of CWD infection. KI mice expressing wild-type deer PrP were subcutaneously vaccinated with recombinant dimeric deer PrP (self-situation), monomeric mouse PrP (non-self), or treated with the adjuvant CpG alone. After several booster injections, mice were infected intraperitoneally with a standard dose of KI mouse-passaged reindeer CWD, without any further boosting. In line with our previous studies in a transgenic mouse model of CWD infection ^35^, we observed humoral immune responses against the immunogens used and an extension of survival time to clinical disease, which is also the experimental endpoint of these studies. Effects on survival were less pronounced than previously reported (28-69 days *vs.* 26-33 days here), but the transgenic mice were over-expressing elk PrP, had a much faster disease progression and earlier experimental endpoints. In addition, in the current study there was no vaccine boosting after CWD challenge, with KI mice surviving for 460 and more days after receiving the last booster dose. When we tested pre-clinical mice at comparable time points, we found lower amounts of PrP^Sc^ in both spinal cord and brain tissues, when compared to the CpG only group. Since mice were infected by a peripheral route, we could observe higher PrP^Sc^ signals in spinal cord tissues, up to around 400 dpi, likely reflection the process of anterograde neuroinvasion. In both vaccine groups some mice were still negative in brain, whereas all 6 tested CpG mice had brain signals and more homogenous signals in the spinal cord. These data show that vaccination targeting PrP^C^ blocks the conversion of PrP^C^ to PrP^Sc^ outside the brain, and delays CWD neuro-invasion.

We next were interested whether vaccination also affects extra-CNS prion replication and the anterograde transport of CWD prions from the CNS to peripheral sites of the body, an aspect never analyzed before. Reduction of shedding of CWD prions from infected animals by vaccination is a key requirement for making vaccines that are able to contain CWD in the long term. The process of prion lateralization is not well understood, and it is unclear whether transport is occurring of PrP^Sc^ preformed in the CNS, or whether generation of newly produced prions outside the brain is involved, or both. The latter would make this process a likely target for vaccine-induced anti-PrP antibodies targeting PrP^C^ mostly. To test whether vaccine-generated antibodies affect CWD prion shedding and likely block prion propagation at peripheral sites of the body, we examined the amounts of PrP^Sc^ in feces and urine of CWD-infected and previously vaccinated KI mice, in comparison to CpG control mice, using ultra-sensitive prion amplification methods. To evade the inhibitory effects of fecal debris and false positive results that were shown in previous studies ^39,45,60^, we combined IOME extraction with PMCA followed by RT-QuIC to increase the specificity of the assay (named here “IPR technique”). IOME was included because it has been shown in previous studies that it enables the detection of low concentrations of PrP^Sc^ in biological samples^42,61,62^.

The new finding of our study is that vaccination with both vaccines (Ddi and Mmo) reduced the shedding of CWD prions in feces and urine. For fecal samples, both the number of positive samples and the amount of seeding activity in samples still testing positive were reduced, when compared to samples taken from CpG-only treated mice. We analyzed fecal samples taken at 200, 250, 300 and 350 dpi, representing about 40%-70% of the incubation time to clinical disease in this model. For assessing prion conversion activity in RT-QuIC, the final readout in IPR, we used established criteria like end-point dilution, time to threshold, maximum of range and area under the curve. Consistently, samples of Ddi or Mmo-vaccinated mice scored significantly lower in these criteria, indicating markedly less prion conversion activity and less CWD prions in these samples when compared to CpG control mice. Overall, best effects were found for Ddi-vaccinated mice, which aligns with our earlier studies that showed the self-antibodies produced by Ddi have a stronger anti-prion effect in cell culture neutralization than Mmo sera (46.2% vs. 5.3%), although titers in ELISA were higher for Mmo-vaccinated mice and cervids ^34^. Yet, the response in the Ddi vaccinated mice seemed to correlate to ELISA titers to some extent. The sera of Ddi mice #7 and 8 that revealed the highest antibody titers showed shedding activity only at one of the four time points analyzed, in addition to low PrP^Sc^ levels in spinal cord.

Of note shedding in feces in CWD-infected KI mice was detectable as early as 200 dpi, the earliest time point we had analyzed here. This matches findings in large animal models, in one elk study as early as 14 dpi ^45^, and 6 months post-infection in white-tailed deer, elk, and mule deer studies ^39,40,43^. Shedding was very consistent throughout testing in the asymptomatic stages in the CpG control group, which matches previous observations in three cervid species ^40^. Shedding is very critical for environmental CWD transmission and long-term persistence of CWD. Of note, in the Ddi-immunized group mice did not shed at various time points (e.g. mice numbers 1, 5 & 7 at 200-300 dpi) or shed only intermittently (mice numbers 3 & 8). The Ddi vaccine candidate is therefore promising for decreasing the CWD contamination in the environment over time.

Findings in urine were even more remarkable. Here, we tested early and very late time points, 150 and 450 days, respectively, corresponding to 30% and >90% of incubation time, with all mice still preclinical. Although we could test urine samples only when pooled and not from individual animals, our findings overall match what we found for fecal samples. The situation for 150 and 250 dpi was almost identical, with no detectable shedding for samples from both Ddi and Mmo-vaccinated mice, whereas 4 serial dilutions of urine tested positive for CpG-only treated mice. For 450 dpi samples, all were negative for Ddi-vaccinated mice (3 animals), whereas Mmo mouse samples were now positive for 4 dilutions (2 mice), and the CpG-only samples (1 mouse) was positive for even 5 serial dilutions. In summary, no detectable prion conversion activity in samples coming from Ddi-vaccinated mice, Mmo-vaccinated mice shed at 450 dpi, and CpG-only treated mice shed at all analyzed time points, with 4-5 log higher titers in the latter when compared to Ddi mice. The pre-clinical detection of CWD in urine aligned with our earlier findings in deer, at 13 and 16 months post infection (mpi) ^63^, and others, at 3 mpi in deer (11) and 18 mpi in white-tailed deer ^43^ until euthanasia at 66 mpi. Several studies in cervids demonstrated that CWD prion concentrations are higher and more consistently detectable in feces than in urine ^11,41,43,60,64^. We observed high prion concentration in both feces and urine samples, using a KI mouse model of CWD infection. Pooled urine samples from the CpG control group showed seeding activity until the 10^-5^ dilution at 450 dpi (**Fig. 5e**), and down to the 10^-7^ dilution at 350 dpi in pooled fecal samples (**Fig 3**), which indicates rather high concentrations of CWD in feces and urine in this KI mouse model, and that the IPR technology described here is very sensitive. Although others had mentioned that it is not necessary to do PMCA plus RT-QuIC after IOME extraction for urine samples of cervids ^43^, this was not the case for our study. For example, IOME with RT-QuIC detected prion seeding activity in pooled urine of the CpG control group only, but when accompanied by PMCA, it detected seeding activity in samples from both the Mmo and CpG groups (**Extended Data Fig. 7**), indicating a higher sensitivity. This shows that the IPR technique described here can be used as a non-invasive method to detect CWD ante-mortem in both feces and urine samples.

In summary, this is the first study to test the efficacy of active vaccination on prion shedding. Our findings in a relevant knock-in mouse model of CWD infection are promising, showing that our vaccine candidates decrease the shedding of CWD prions in urine and feces. Whether this has the potential to break the CWD transmission cycle in the long term needs to be determined in vaccine studies in deer and elk, some of them ongoing. Taken together, the vaccination strategy described here has obviously two additive effects. It has the potential to improve individual survival, which will hopefully translate into population effects. Importantly, it also reduces prion shedding, likely translating into reduction of CWD prions in the environment in the long term.

## Materials & Methods

### Ethical statement

Four- to eight-week-old female KI mice expressing wild-type cervid PrP were used ^37^. All animal experiments were approved by the University of Calgary Health Sciences Animal Care Committee (protocol# AC18-0138 and AC22-0106), following the guidelines issued by the Canadian Council on Animal Care.

### CWD prion material

Brain material from a CWD-infected captive white-tailed deer (WTD) was used to orally infect reindeer experimentally ^65^. This reindeer developed clinical signs of prion infection, and the brain was positive in WB and ELISA. Brain homogenate (BH) of the infected reindeer was used to challenge KI mice intraperitoneally ^37^, leading to a 100% attack rate. We used this KI mouse-adapted reindeer CWD in our study. 1% BH in PBS was used as inoculum.

### Mouse bioassay

We used three groups of mice (n=8 per group): Ddi or Mmo vaccinated groups and a CpG control group. The first bleeding was done immediately before starting the vaccination procedure and used as pre-immune sera. The vaccine groups were subcutaneously (s.c.) injected with a priming dose of 100 μg of protein plus 5 μM CpG as adjuvant, followed by four booster doses consisting of 50 μg protein plus 5 μM CpG. The control group received 5 μM CpG in PBS only. Post-immune sera were collected 10 days after the fourth booster dose, then all groups were challenged intraperitoneally (i.p.) with 1% BH KI mouse-adapted reindeer CWD **(Fig. 1a)**.

### Feces and urine collection and processing

Individual or pooled feces and pooled urine were collected every 50 days post-inoculation (dpi) **(Fig. 1b)**. Feces were homogenized in sterile 1× PBS 10% (w/v) homogenates in FastPrep-24™ Lysing D Matrix tubes with ceramic beads (MP Biomedicals) using the FastPrep-24™ Tissue Homogenizer (MP Biomedicals). Fecal homogenates were centrifuged twice at 18,000 x g for 10 minutes to remove debris and were aliquoted and stored at −80°C until further use. Superparamagnetic iron oxide beads (IOME) [≈9μm; BM547 Bangs Laboratories, Indiana] were used to concentrate PrP^Sc^ as previously described ^42^. Briefly, 2 μl of IOME beads were washed once with 500 μl PBS using a magnetic particle separator (MPS) (Pure Biotech, New Jersey), and supernatant was removed while the tubes were in the MPS. 500 μl of 10% fecal homogenate were added, and samples were placed on the end-over-end rocker for 2 hours at room temperature. Then, tubes were placed in MPS for 5 minutes, supernatant discarded, and beads were resuspended in 10 μl of 0.1 % sodium dodecyl sulphate (SDS) in PBS and samples stored at −80°C until further use.

### Protein misfolding cyclic amplification assay (PMCA)

PMCA was performed using the protocol published by Arifin et al. ^37^ with some modifications. PMCA conversion buffer was prepared, which contains 4 mM EDTA, 1% Triton X-100 and 1 tablet cOmplete™ Protease Inhibitor Mini (Roche) in 1 x PBS. For the PMCA substrate, in a prion-free area, brains from naïve KI mice were collected and homogenized as 10% (w/v) BH using a Potter–Elvehjem PTFE pestle and glass tube (Sigma-Aldrich, #P7984) in ice-cold PMCA conversion buffer. The brain homogenates were aliquoted into 500 μL in sterile Eppendorf tubes and stored at −80 °C until use. In 0.2 mL tubes (ThermoFisher, #AB0337), three PTFE balls (McMaster-Carr, #9660K12, 3/32 in ø) and 90 μL of substrate were added. Moving to the prion area, the water bath (CC304-B, Huber) was set at 36.5 °C. The extracted 10% fecal homogenate or urine sample was diluted 1:10 in PMCA conversion buffer. Ten μL of each seed dilution was added to the 0.2 mL tubes, and all reactions were conducted in duplicate. Finally, PMCA tubes were sealed with parafilm, decontaminated in a 2.5 % bleach for 5 min, rinsed with H_2_O, and placed in a tube rack inside a microplate sonicator horn (431MPXH and Q700, QSonica) connected to the circulating water bath. Each cycle of PMCA is conducted for 24 hours which equals 48 cycles of 30 s sonication at 375 to 395 W followed by 29.5 min rest. For serial PMCA (sPMCA), 10 μL of PMCA product from the first round was transferred into 90 μL of fresh PMCA substrate and run for another 24 h with the same settings. This process was repeated to obtain three rounds of sPMCA with a total of 144 cycles. The PMCA products of the third round were used as a seed in the RT-QuIC assay.

### Real-time quaking induced conversion (RT-QuIC) assay

As readout of the PMCA products for fecal homogenates after IOME RT-QuIC was performed as previously described, using recombinant mouse PrP (aa23-231) as the substrate ^63^. The reaction mixture included 10 μg of recombinant PrP, 10 μM Thioflavin-T (ThT), 170 mM NaCl, 1× PBS (containing 130 mM NaCl), 1 mM EDTA, and water. Each well of a 96-well plate (Millipore Sigma, Ref# 3603) received 98 μL of this mixture plus 2 μL of the seed (PMCA products of 10% fecal homogenate extracted using IOME), tenfold serially diluted in RT-QuIC seed dilution buffer in quadruplicate. Plates were sealed, incubated at 42°C, and shaken at 700 rpm in a BMG Labtech FLUOstar™ plate reader for 50 hours. Thioflavin T fluorescence was measured every 15 minutes. A sample was considered positive if two out of the four replicates exceeded the threshold, defined as average RFU of the negative control plus five standard deviations.

### Preparation of the immunogen

Recombinant monomeric mouse (Mmo) and dimeric deer (Ddi) prion proteins were prepared as described previously ^31,34^. Mmo consists of N-terminally his-tagged PrP encompassing amino acids (aa) 23–231 of mouse PrP, not containing the N- and C-terminal peptides (aa 1–22 and 232– 254, respectively). The Ddi encoding constructs were synthesized by GeneArt and include two sequences of aa 23-231 of deer PrP, linked together by a short 7-amino acid linker (AGAIGGA). In brief, all constructs were cloned into the pQE30 expression vector (Qiagen) and expressed in *E. coli* BL21-Gold (DE3) pLysS cells (Stratagene). Guanidine hydrochloride 6M (pH 7.8) was used to lyse the bacterial cells and centrifuged at 6,000 x g for 30 minutes to remove cell debris. Protein purification was done through Ni-affinity chromatography using Ni-NTA Superflow resin (Qiagen). The purified protein was eluted with the elution buffer and refolded using the dialysis buffer. Quality control tests were done on the produced protein, which included the BCA protein assay kit (Pierce, Thermo Scientific) to measure protein concentration, Coomassie blue staining of SDS-PAGE, Western blot and ELISA.

### Proteinase-K (PK) digestion

Brain homogenates (20%) or spinal cord homogenate (10%) prepared in 1X PBS were mixed with an equal volume of 2X lysis buffer and were digested with 50 μg/ml of PK at 37°C for one hour. Aliquots of the PMCA products of the feces and urine samples were mixed with 2x lysis buffer and digested with 50 μg/ml and 37 μg/ml of PK, respectively. Pefabloc protease inhibitor (1x; VWR) was added to stop the enzymatic reaction, followed by adding 3 x SDS sample loading buffer and boiling for 10 minutes at 95°C.

### SDS-PAGE and Western blot

Samples were separated on a 12.5% SDS-poly-acrylamide gel and transferred to PVDF membranes (Amersham, GE Healthcare). Membranes were blocked with 5% non-fat milk in Tris-buffered saline with a final concentration of 0.1% Tween-20 (TBST) for 1 hr at room temperature. Membranes were probed with the anti-PrP monoclonal antibody 4H11 (1:500 or 1:700) ^66^ followed by washing with TBST. Horseradish peroxidase-conjugated goat anti-mouse IgG antibody (Sigma) was used as secondary antibody (1:5,000), followed by washing with TBST. Luminita horseradish peroxidase substrate (Millipore Sigma) was used for developing. Images were acquired on X-ray films (Denville Scientific). Image J software was used to quantify and determine the relative values of PrP^Sc^ signals. Calculations were done on Microsoft Excel, and graphs generated using GraphPad Prism 10.

### ELISA

ELISA was conducted as described previously ^34^. In brief, 1 μg of either Ddi or Mmo recombinant protein in sodium-carbonate buffer (pH 9.5) was used to coat high-binding 96-well plates (Greiner Bio-One GmbH, Frickenhausen, Germany) overnight at room temperature. Plates were washed with PBS-T, then blocked using 3% bovine serum albumin (BSA) in PBS-T for 2 h at 37 °C. The post-immune sera from every mouse, diluted 1:100 or serially diluted in 3% BSA, were added to the plates, incubated for 1 h, and plates washed with PBS-T. The secondary antibody used was HRP-labeled anti-mouse IgG antibody (Jackson ImmunoResearch, West Grove, PA) diluted 1:4,000 in 3% BSA. Signal detection was performed using the ABTS peroxidase substrate system (Kirkegaard & Perry Laboratories). To assess the ELISA signal, the optical density (OD) was measured at 405 nm using a BioTek Synergy HT microplate reader.

### Epitope mapping

We tested 14 peptides covering the full-length mature cervid PrP, with each peptide consisting of 20 aa peptides with 5 aa overlap **(Extended Data Table 1)**. Peptide 6a contains the 3F4 epitope, while peptide 6b represents the corresponding murine sequence. We used the same protocol as described by us previously ^31,34^. Using CovaLink NH microtiter plates, activation of the wells was done using DSS bifunctional linker in carbonate buffer. Coating was done with 10 μg of each peptide or 1 μg of recombinant protein (control) at room temperature overnight. After washing, wells were blocked with BSA for 30 minutes at 37 °C and incubated with post-immune sera (1:100 in blocking buffer) for 2 hr at 37 °C. Plate were washed and incubated with secondary antibodies (1:4,000 in blocking buffer) for 1hr at 37 °C. After final washing, wells were incubated with ABTS substrate, the optical density was measured at 405 nm.

### Statistics

Statistical analyses were done using GraphPad Prism software (GraphPad 10.4.1, Software, USA). For statistical analysis of immunoblot signals, two-tailed independent *Student’s t-test* groups or one-way analysis of *variance (ANOVA)*, followed by *Turkey’s multiple comparison test*, as applicable, were used. The area under the curve (AUC), time to threshold, and maximum of range were calculated using Omega data analysis (MARS version 4.01 R2). Statistical significance was tested using one-way analysis of variance (*ANOVA*) followed by *Turkey’s multiple comparison test.* Values are expressed as mean ± SEM. Significance = \**p ≤ 0.05*, \*\**p ≤ 0.01*, and \*\*\**p ≤ 0.001*.

## Author contributions

HMS and HAH designed the study, wrote the manuscript, and organized the data. HAH conducted the animal bioassay, processing samples, IOME, PMCA, RT-QuIC, immunoblotting, and data analysis. DA contributed to mouse bioassay, protein preparation, ELISA and epitope mapping. GC conducted to the animal bioassay and protein preparation for RT-QuIC, KL assisted with sample collection and SP contributed to the ELISA end point dilution assay. CD and BK participated in protein preparation. SG provided the KI mice and shared the inocula used in this study. LK and WSJ helped to generate the KI mice. All authors reviewed and approved the manuscript for publication. HMS is the corresponding author and assumes full responsibility for the manuscript.

## Materials & Correspondence

Supplementary Information is available for this paper. Correspondence and material requests should be addressed to Dr. Hermann Schatzl.

## List of Supplementary Materials

Supplementary Information

SFig1

SFig2

SFig3

SFig4

SFig5

SFig6

SFig7

Table S1

Table S2

Table S3

## References

1. Prusiner, S. B. Novel proteinaceous infectious particles cause scrapie. Science. 1982; 216, 136–144, doi:10.1126/science.6801762.

2. Prusiner, S. B. Prions. Proc Natl Acad Sci USA. 1998;95, 13363–13383, doi:10.1073/pnas.95.23.13363.

3. USGS. Expanding Distribution of Chronic Wasting Disease, 2024; <https://www.usgs.gov/centers/nwhc/science/expanding-distribution-chronic-wasting-disease>.

4. Williams, E. S. & Young, S. Chronic wasting disease of captive mule deer: a spongiform encephalopathy. J Wildl Dis 1980;16, 89–98, doi:10.7589/0090-3558-16.1.89.

5. Napper, S. & Schatzl, H. M. Vaccines for prion diseases: a realistic goal? Cell Tissue Res. 2023;392, 367–392, doi:10.1007/s00441-023-03749-7.

6. Tranulis, M. A. et al. Chronic wasting disease in Europe: new strains on the horizon. Acta Vet Scand. 2021;63, 48, doi:10.1186/s13028-021-00606-x.

7. Otero, A. et al. Prion protein polymorphisms associated with reduced CWD susceptibility limit peripheral PrP(CWD) deposition in orally infected white-tailed deer. BMC Vet Res. 2019;15, 50, doi:10.1186/s12917-019-1794-z.

8. Fox, K. A., Jewell, J. E., Williams, E. S. & Miller, M. W. Patterns of PrPCWD accumulation during the course of chronic wasting disease infection in orally inoculated mule deer (Odocoileus hemionus). J Gen Virol. 2006;87, 3451–3461, doi:10.1099/vir.0.81999-0.

9. Sigurdson, C. J. & Aguzzi, A. Chronic wasting disease. Biochim Biophys Acta. 2007;1772, 610–618, doi:10.1016/j.bbadis.2006.10.010.

10. Haley, N. J. et al. Detection of chronic wasting disease prions in salivary, urinary, and intestinal tissues of deer: potential mechanisms of prion shedding and transmission. J Virol. 2011;85, 6309–6318, doi:10.1128/JVI.00425-11.

11. Henderson, D. M. et al. Longitudinal Detection of Prion Shedding in Saliva and Urine by Chronic Wasting Disease-Infected Deer by Real-Time Quaking-Induced Conversion. J Virol. 2015;89, 9338–9347, doi:10.1128/JVI.01118-15.

12. Mathiason, C. K. et al. Infectious prions in pre-clinical deer and transmission of chronic wasting disease solely by environmental exposure. PLoS One. 2009;4, e5916, doi:10.1371/journal.pone.0005916.

13. Pritzkow, S. Transmission, Strain Diversity, and Zoonotic Potential of Chronic Wasting Disease. Viruses. 2022;14, doi:10.3390/v14071390.

14. Hamir, A. N., Miller, J. M., Kunkle, R. A., Hall, S. M. & Richt, J. A. Susceptibility of cattle to first-passage intracerebral inoculation with chronic wasting disease agent from white-tailed deer. Vet Pathol. 2007;44, 487–493, doi:10.1354/vp.44-4-487.

15. Hamir, A. N. et al. Experimental transmission of chronic wasting disease agent from mule deer to cattle by the intracerebral route. J Vet Diagn Invest. 2005;17, 276–281, doi:10.1177/104063870501700313.

16. Hamir, A. N. et al. Transmission of chronic wasting disease of mule deer to Suffolk sheep following intracerebral inoculation. J Vet Diagn Invest. 2006;18, 558–565, doi:10.1177/104063870601800606.

17. Heisey, D. M. et al. Chronic wasting disease (CWD) susceptibility of several North American rodents that are sympatric with cervid CWD epidemics. J Virol. 2010;84, 210–215, doi:10.1128/JVI.00560-09.

18. Raymond, G. J. et al. Transmission and adaptation of chronic wasting disease to hamsters and transgenic mice: evidence for strains. J Virol. 2007;81, 4305–4314, doi:10.1128/JVI.02474-06.

19. Race, B. et al. Susceptibilities of nonhuman primates to chronic wasting disease. Emerg Infect Dis. 2009;15, 1366–1376, doi:10.3201/eid1509.090253.

20. Hannaoui, S. et al. Transmission of cervid prions to humanized mice demonstrates the zoonotic potential of CWD. Acta Neuropathol. 2022;144, 767–784, doi:10.1007/s00401-022-02482-9.

21. Race, B. et al. Lack of Transmission of Chronic Wasting Disease to Cynomolgus Macaques. J Virol. 2018;92, doi:10.1128/JVI.00550-18.

22. Race, B. et al. Chronic wasting disease agents in nonhuman primates. Emerg Infect Dis. 2014;20, 833–837, doi:10.3201/eid2005.130778.

23. Marsh, R. F., Kincaid, A. E., Bessen, R. A. & Bartz, J. C. Interspecies transmission of chronic wasting disease prions to squirrel monkeys (Saimiri sciureus). J Virol. 2005;79, 13794–13796, doi:10.1128/JVI.79.21.13794-13796.2005.

24. Groveman, B. R. et al. Lack of Transmission of Chronic Wasting Disease Prions to Human Cerebral Organoids. Emerg Infect Dis. 2024;30, 1193–1202, doi:10.3201/eid3006.231568.

25. Angers, R. C. et al. Prions in skeletal muscles of deer with chronic wasting disease. Science. 2006;311, 1117, doi:10.1126/science.1122864.

26. Angers, R. C. et al. Chronic wasting disease prions in elk antler velvet. Emerg Infect Dis. 2009;15, 696–703, doi:10.3201/eid1505.081458.

27. Schwarz, A. et al. Immunisation with a synthetic prion protein-derived peptide prolongs survival times of mice orally exposed to the scrapie agent. Neurosci Lett. 2003;350, 187–189, doi:10.1016/s0304-3940(03)00907-8.

28. Sigurdsson, E. M. et al. Anti-prion antibodies for prophylaxis following prion exposure in mice. Neurosci Lett. 2003;336, 185–187, doi:10.1016/s0304-3940(02)01192-8.

29. White, A. R. et al. Monoclonal antibodies inhibit prion replication and delay the development of prion disease. Nature. 2003;422, 80–83, doi:10.1038/nature01457.

30. Heppner, F. L. et al. Prevention of scrapie pathogenesis by transgenic expression of anti-prion protein antibodies. Science. 2001;294, 178–182, doi:10.1126/science.1063093.

31. Gilch, S. et al. Polyclonal anti-PrP auto-antibodies induced with dimeric PrP interfere efficiently with PrPSc propagation in prion-infected cells. J Biol Chem. 2003;278, 18524–18531, doi:10.1074/jbc.M210723200.

32. Polymenidou, M. et al. Humoral immune response to native eukaryotic prion protein correlates with anti-prion protection. Proc Natl Acad Sci USA. 2004;101 Suppl 2, 14670–14676, 29 doi:10.1073/pnas.0404772101.

33. Kaiser-Schulz, G. et al. Polylactide-coglycolide microspheres co-encapsulating recombinant tandem prion protein with CpG-oligonucleotide break self-tolerance to prion protein in wild-type mice and induce CD4 and CD8 T cell responses. J Immunol. 2007;179, 2797–2807, doi:10.4049/jimmunol.179.5.2797.

34. Abdelaziz, D. H. et al. Immunization of cervidized transgenic mice with multimeric deer prion protein induces self-antibodies that antagonize chronic wasting disease infectivity in vitro. Sci Rep. 2017;7, 10538, doi:10.1038/s41598-017-11235-8.

35. Abdelaziz, D. H. et al. Recombinant prion protein vaccination of transgenic elk PrP mice and reindeer overcomes self-tolerance and protects mice against chronic wasting disease. J Biol Chem. 2018;293, 19812–19822, doi:10.1074/jbc.RA118.004810.

36. Napper, S. & Schatzl, H. M. Oral vaccination as a potential strategy to manage chronic wasting disease in wild cervid populations. Front Immunol. 2023;14, 1156451, doi:10.3389/fimmu.2023.1156451.

37. Arifin, M. I. et al. Heterozygosity for cervid S138N polymorphism results in subclinical CWD in gene-targeted mice and progressive inhibition of prion conversion. Proc Natl Acad Sci USA. 2023;120, e2221060120, doi:10.1073/pnas.2221060120.

38. Pulford, B. et al. Detection of PrPCWD in feces from naturally exposed Rocky Mountain elk (Cervus elaphus nelsoni) using protein misfolding cyclic amplification. J Wildl Dis. 2012;48, 425–434, doi:10.7589/0090-3558-48.2.425.

39. Hwang, S., Greenlee, J. J. & Nicholson, E. M. Real-Time Quaking-Induced Conversion Detection of PrP(Sc) in Fecal Samples From Chronic Wasting Disease Infected White-Tailed Deer Using Bank Vole Substrate. Front Vet Sci. 2021;8, 643754, doi:10.3389/fvets.2021.643754.

40. Plummer, I. H., Wright, S. D., Johnson, C. J., Pedersen, J. A. & Samuel, M. D. Temporal patterns of chronic wasting disease prion excretion in three cervid species. J Gen Virol. 2017;98, 1932–1942, doi:10.1099/jgv.0.000845.

41. Tennant, J. M. et al. Shedding and stability of CWD prion seeding activity in cervid feces. PLoS One. 2020;15, e0227094, doi:10.1371/journal.pone.0227094.

42. Denkers, N. D., Henderson, D. M., Mathiason, C. K. & Hoover, E. A. Enhanced prion detection in biological samples by magnetic particle extraction and real-time quaking-induced conversion. J Gen Virol. 2016;97, 2023–2029, doi:10.1099/jgv.0.000515.

43. Denkers, N. D. et al. Temporal Characterization of Prion Shedding in Secreta of White-Tailed Deer in Longitudinal Study of Chronic Wasting Disease, United States. Emerg Infect Dis. 2024;30, 2118–2127, doi:10.3201/eid3010.240159.

44. Arifin, M. I. et al. Cervid Prion Protein Polymorphisms: Role in Chronic Wasting Disease Pathogenesis. Int J Mol Sci. 2021;22, doi:10.3390/ijms22052271.

45. Cheng, Y. C. et al. Early and Non-Invasive Detection of Chronic Wasting Disease Prions in Elk Feces by Real-Time Quaking Induced Conversion. PLoS One. 2016;11, e0166187, doi:10.1371/journal.pone.0166187.

46. Pritzkow, S. et al. Grass plants bind, retain, uptake, and transport infectious prions. Cell Rep. 2015;11, 1168–1175, doi:10.1016/j.celrep.2015.04.036.

47. Park, K. J. et al. Detection of chronic wasting disease prions in the farm soil of the Republic of Korea. mSphere. 2025;10, e0086624, doi:10.1128/msphere.00866-24.

48. Kuznetsova, A. et al. Detection of Chronic Wasting Disease Prions in Prairie Soils from Endemic Regions. Environ Sci Technol. 2024;58, 10932–10940, doi:10.1021/acs.est.4c04633.

49. Beekes, M. & McBride, P. A. The spread of prions through the body in naturally acquired transmissible spongiform encephalopathies. FEBS J. 2007;274, 588–605, doi:10.1111/j.1742-4658.2007.05631.x.

50. Wisniewski, T. & Goni, F. Immunomodulation for prion and prion-related diseases. Expert Rev Vaccines. 2010;9, 1441–1452, doi:10.1586/erv.10.131.

51. Aucouturier, P., Carp, R. I., Carnaud, C. & Wisniewski, T. Prion diseases and the immune system. Clin Immunol. 2000;96, 79–85, doi:10.1006/clim.2000.4875.

52. Taschuk, R. et al. Induction of PrP(Sc)-specific systemic and mucosal immune responses in white-tailed deer with an oral vaccine for chronic wasting disease. Prion. 2017;11, 368–380, doi:10.1080/19336896.2017.1367083.

53. Taschuk, R. et al. Safety, specificity and immunogenicity of a PrP(Sc)-specific prion vaccine based on the YYR disease specific epitope. Prion. 2014;8, 51–59, doi:10.4161/pri.27962.

54. Wood, M. E. et al. Accelerated onset of chronic wasting disease in elk (Cervus canadensis) vaccinated with a PrP(Sc)-specific vaccine and housed in a prion contaminated environment. Vaccine. 2018;36, 7737–7743, doi:10.1016/j.vaccine.2018.10.057.

55. Fleming M, F. A., Tancowny B, Telling G, Wille H. Optimizing prion vaccination in a transgenic mouse model of Gerstmann-Sträussler-Scheinker disease. Prion. 2022;16, 95–253, doi:10.1080/19336896.2022.2091286.

56. Bueler, H. et al. Normal development and behaviour of mice lacking the neuronal cell-surface PrP protein. Nature. 1992;356, 577–582, doi:10.1038/356577a0.

57. Mallucci, G. R. et al. Post-natal knockout of prion protein alters hippocampal CA1 properties, but does not result in neurodegeneration. EMBO J. 2002;21, 202–210, doi:10.1093/emboj/21.3.202.

58. Seelig, D. M., Mason, G. L., Telling, G. C. & Hoover, E. A. Pathogenesis of chronic wasting disease in cervidized transgenic mice. Am J Pathol. 2010;176, 2785–2797, doi:10.2353/ajpath.2010.090710.

59. Bian, J. et al. Primary structural differences at residue 226 of deer and elk PrP dictate selection of distinct CWD prion strains in gene-targeted mice. Proc Natl Acad Sci USA. 2019;116, 12478–12487, doi:10.1073/pnas.1903947116.

60. Henderson, D. M. et al. Detection of chronic wasting disease prion seeding activity in deer and elk feces by real-time quaking-induced conversion. J Gen Virol. 2017;98, 1953–1962, doi:10.1099/jgv.0.000844.

61. Miller, M. B. & Supattapone, S. Superparamagnetic nanoparticle capture of prions for amplification. J Virol. 2011;85, 2813–2817, doi:10.1128/JVI.02451-10.

62. Kraft, C. N., Denkers, N. D., Mathiason, C. K. & Hoover, E. A. Longitudinal detection of prion shedding in nasal secretions of CWD-infected white-tailed deer. J Gen Virol. 2023;104, doi:10.1099/jgv.0.001825.

63. John, T. R., Schatzl, H. M. & Gilch, S. Early detection of chronic wasting disease prions in urine of pre-symptomatic deer by real-time quaking-induced conversion assay. Prion. 2013;7, 253–258, doi:10.4161/pri.24430.

64. Haley, N. J., Seelig, D. M., Zabel, M. D., Telling, G. C. & Hoover, E. A. Detection of CWD prions in urine and saliva of deer by transgenic mouse bioassay. PLoS One. 2009;4, e4848, doi:10.1371/journal.pone.0004848.

65. Mitchell, G. B. et al. Experimental oral transmission of chronic wasting disease to reindeer (*Rangifer tarandus tarandus*). PLoS One 2012;7, e39055, doi:10.1371/journal.pone.0039055.

66. Ertmer, A. et al. The tyrosine kinase inhibitor STI571 induces cellular clearance of PrP^Sc^ in prion-infected cells. J. Biol. Chem. 2004;279, 41918–41927, doi:10.1074/jbc.M405652200.

