## Supplementary data for "Prion shedding is reduced by chronic wasting disease vaccination"

### Extended Data

#### Extended Data Fig. 1

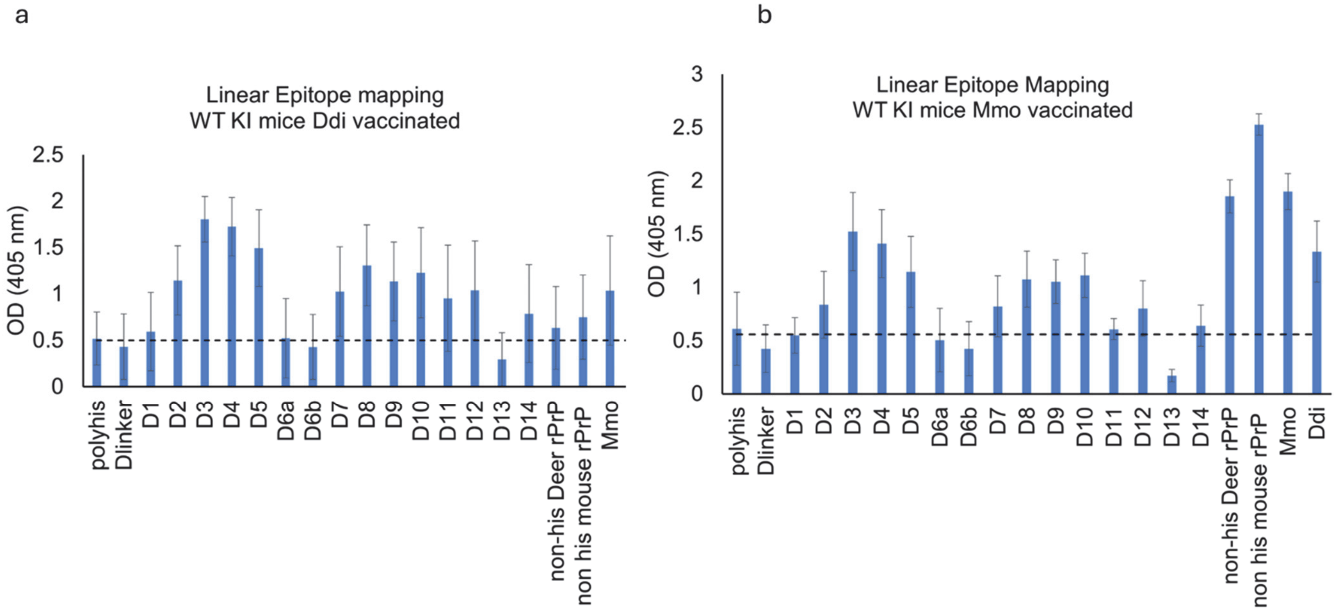


**Extended Data Fig. 1. Linear epitope mapping for Ddi and Mmo vaccinated mice.** The Y-axis represents the optical density (OD) at 405 nm, indicating the reactivity of the post-immune sera from either **a,** Ddi or **b,** Mmo immunized mice against the x-axis represented by the linear epitopes of deer rPrP, as measured by ELISA. The dashed horizontal line represents the cut-off, which is 3 times the average of pre-immune sera. The data in panel **a**, represents the average of Ddi sera 1,3,4 and 6. The data in panel **b,** represents the average of Mmo sera numbers 1,2,3 and 6. Data are presented as mean ± SD of results from four individual mice of each group. The amino acid sequences for linear epitopes are shown in Extended data table 1.

#### Extended Data Fig. 2

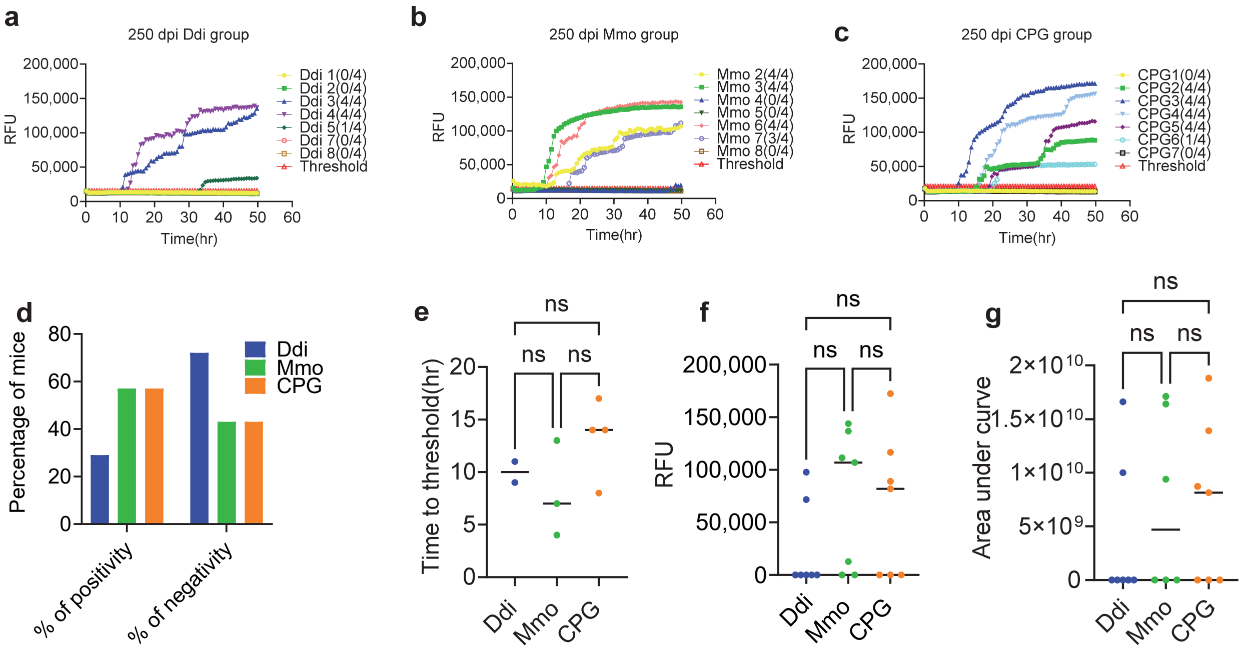


**Extended Data fig. 2. RT-QuIC data showing the seeding activity in feces from vaccinated and control group at 250 dpi.**10% individual fecal homogenate from all groups extracted using IOME followed by three rounds of PMCA reactions using KI $PrP^{C}$ substrate seeded with $10^{-1}$ dilution of 250 dpi 10% fecal homogenate, in duplicates for all vaccinated and control group. Positive control for the PMCA was naïve feces spiked with mouse-adapted CWD reindeer and negative control for the PMCA was naïve feces. All samples and controls were subjected to PMCA reaction. The PMCA products analyzed using RT-QuIC assay at $10^{-2}$ dilution. **a-c,** Representative RT-QuIC graphs showing the seeding activity in feces from vaccinated or control KI mice. Samples were considered positive when 2 out of 4 wells crossed the threshold, which defined as the average RFU of the negative control group plus five times its standard deviation. The y-axis represents the RFU, and the x-axis represents the time in hours(hr). **d,** Chi square test, **e,** Time to threshold, **f,** Maximum of range**, g,** Area under curve. Graphs were generated using GraphPad Prism (version 10). Statistical analysis done using Chi-square test and **** P-value< 0.0001- or One-way ANOVA followed by a Tukey's multiple comparison and ns: not significant.

#### Extended Data Fig. 3

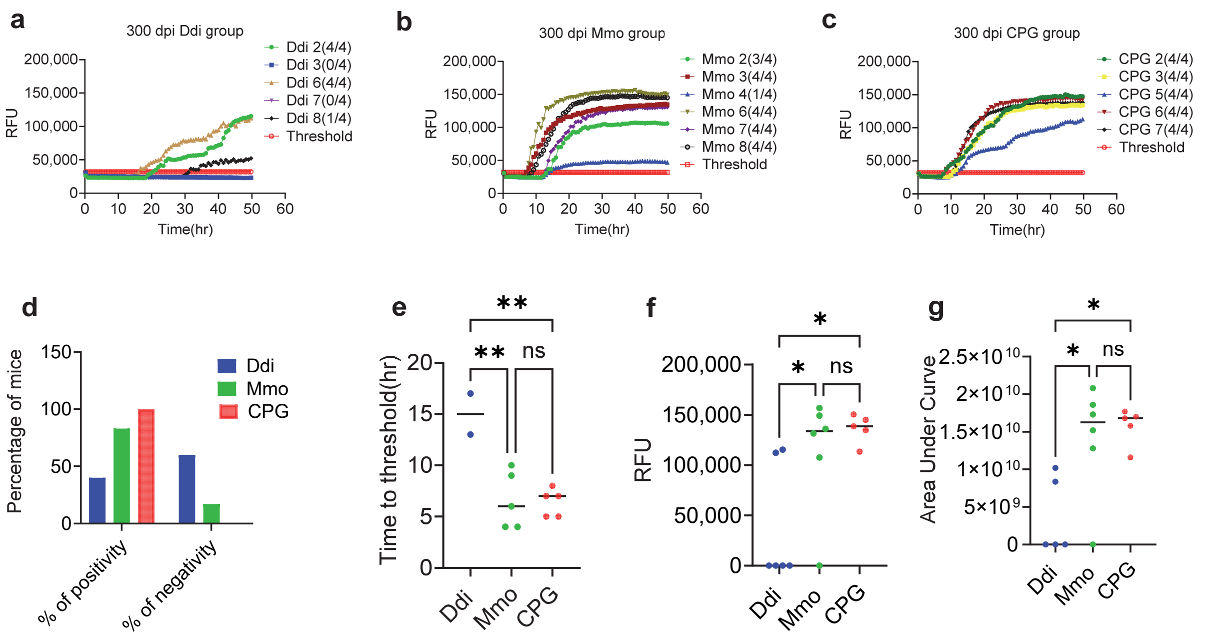


**Extended Data Fig. 3. RT-QuIC analysis of the PMCA products illustrating the difference in the CWD shedding between KI vaccinated and control group at 300 dpi.** First,10% individual fecal homogenate from all groups extracted using IOME. Then, KI $PrP^C$ substrate seeded with $10^{-1}$ dilution of the extracted fecal homogenate, in duplicates for all vaccinated and control group followed by three rounds of PMCA reactions. Positive control for the PMCA was naïve feces spiked with mouse-adapted CWD reindeer and negative control for the PMCA was naïve feces. The PMCA products analyzed using RT-QuIC assay at $10^{-1}$ dilution. **a-c,** representative RT-QuIC graphs showing the seeding activity in feces from vaccinated or control KI mice. Samples were considered positive when 2 out of 4 wells crossed the threshold, which defined as the average RFU of the negative control group plus five times its standard deviation. The y-axis represents the RFU, and the x-axis represents the time in hours(hr). **d,** Chi square test, **e,** Time to threshold, **f,** Maximum of range and **g,** Area under curve. Graphs were generated using GraphPad Prism (version 10). Statistical analysis done using Chi-square test and **** P-value< 0.0001 or One-way ANOVA followed by a Tukey's multiple comparison. For time to threshold: Ddi vs. Mmo ** P-value= 0.0043, Ddi vs. CPG ** P-value= 0.0037. For maximum of range: Ddi vs. Mmo * P-value= 0.0498, Ddi vs. CPG * P-value= 0.0149. ns: not significant. For area under curve: Ddi vs. Mmo * P-value= 0.0225, Ddi vs. CPG * P-value= 0. 0120.

#### Extended Data Fig. 4

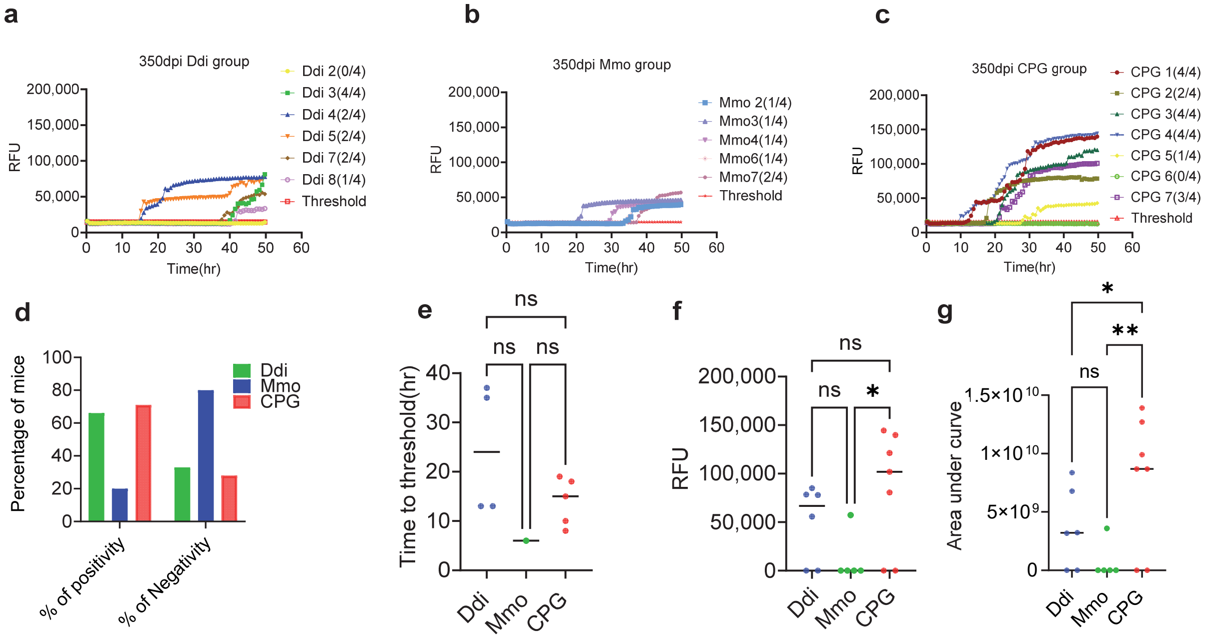


**Extended Data Fig. 4. Readouts of the PMCA showing the difference in the CWD shedding between KI vaccinated and control mice groups at 350 dpi.** KI $PrP^C$ substrate seeded with $10^{-1}$ dilution of the extracted 10% fecal homogenate, in duplicates for all vaccinated and control group followed by three rounds of PMCA reactions. Positive control for the PMCA was naïve feces spiked with mouse-adapted CWD reindeer and negative control for the PMCA was naïve feces. The PMCA products analyzed using RT-QuIC assay at $10^{-5}$ dilution. Panels **a-c,** RT-QuIC graphs showing the seeding activity in feces from vaccinated or control KI mice. Samples were considered positive when 2 out of 4 wells crossed the threshold, which defined as the average RFU of the negative control group plus five times its standard deviation. The y-axis represents the RFU, and the x-axis represents the time in hours(hr). **d,** Chi square test, **e,** Time to threshold, **f,** Maximum of range, **h,** Area under curve. Graphs were generated using GraphPad Prism (version 10). Statistical analysis done using Chi-square test and **** P-value< 0.0001 or Two-way ANOVA followed by a Tukey's multiple comparison for Area under curve Ddi vs CPG * P-value= 0.0453 and Mmo vs CPG ** P-value= 0.0052. For Maximum of range * P-value=0.0308.

**Extended Data Fig. 5**

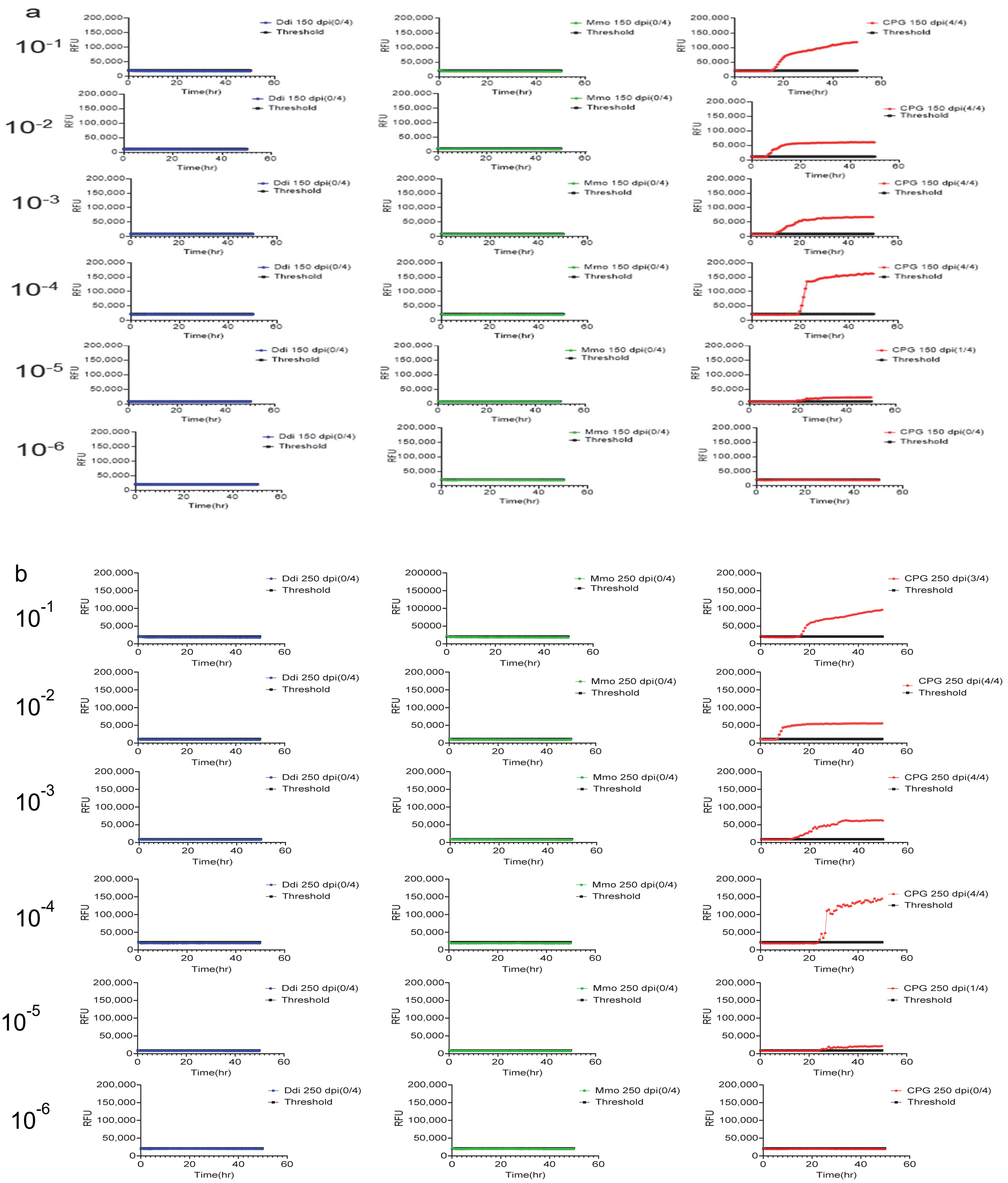

**Extended Data Fig. 5. Prion seeding activity in third round PMCA products for urine of KI mice vaccinated with Ddi or Mmo and control CPG group, all groups were intraperitoneally inoculated with reindeer mouse-adapted CWD.** The graphs represent RT-QuIC results of serially diluted ($10^{-1}$ to $10^{-6}$) for **a,**150 dpi and **b,** 250dpi urine samples from all groups after three rounds of PMCA using mouse rPrP substrate. Fluorescence signals were measured every 15 min. The *x*-axis represents the reaction time (hour), the *y*-axis represents the relative fluorescence units (RFU), and each curve represents a different dilution for a different group. The threshold was based on the average fluorescence values of the negative control + 5 × SD used in every assay. RT-QuIC results from urine with mouse-adapted reindeer CWD were included as positive control, and results from non-inoculated KI urine were included as negative control. Both negative and positive controls are included in the PMCA. Each curve represents an average of 4 technical replicates.

**Extended Data Fig. 6**

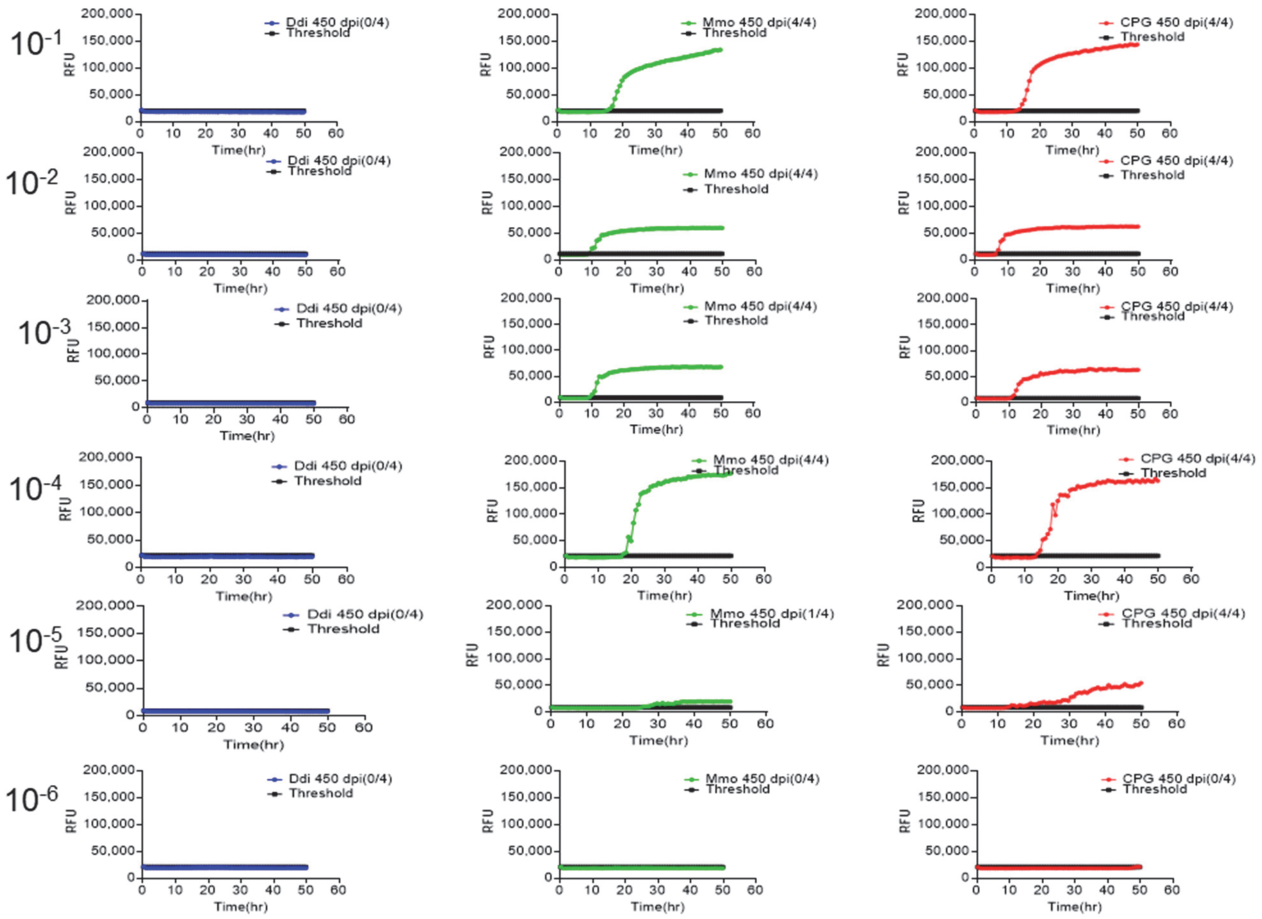


**Extended DataFig.6. RT-QuIC data showing the CWD prion seeding activity in third round PMCA products for urine of KI mice vaccinated and control groups at 450dpi.**All groups were intraperitoneally inoculated with mouse-adapted reindeer CWD. The graphs represent RT-QuIC results of serially diluted ($10^{-1}$ to $10^{-6}$) for 450dpi urine samples from all groups after three rounds of PMCA using mouse rPrP substrate. Fluorescence signals were measured every 15 min. The x-axis represents the reaction time (hours), the y-axis represents the relative fluorescence units (RFU), and each curve represents a different dilution for different groups. The threshold was based on the average fluorescence values of the negative control + 5 × SD used in every assay. RT-QuIC results from urine with mouse-adapted reindeer CWD were included as positive control, and results from non-inoculated KI urine were included as negative control. Both negative and positive controls are included in the PMCA. Each curve represents the average of 4 technical replicates.

#### Extended Data Fig. 7

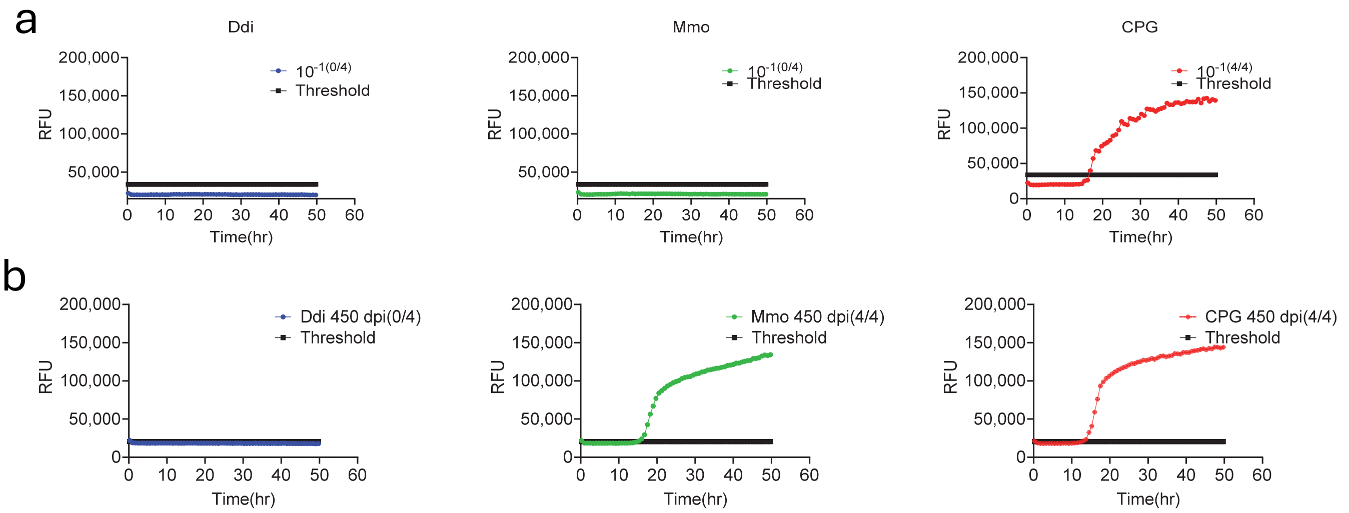


**Extended Data Fig. 7. RT-QuIC data showing the seeding activity in 450dpi pooled urine samples extracted with a, IOME only or b, IOME followed by three rounds of PMCA using mouse rPrP substrate.** Fluorescence signals were measured every 15 min. The *x*-axis represents the reaction time (hours), the *y*-axis represents the relative fluorescence units (RFU), and each curve represents $10^{-1}$ dilution for different groups. The threshold was based on the average fluorescence values of the negative control + 5 × SD used in every assay. RT-QuIC results from urine with mouse-adapted reindeer CWD were included as positive control, and results from non-inoculated KI urine were included as negative control. Both negative and positive controls are included in the PMCA. Each curve represents an average of 4 technical replicates.

**Extended Data Table 1:** Linear epitope sequence for the deer rPrP.

| Epitope | Sequence |
|---|---|
| Polyhistag | MRG SHH HHH HGS CKK RPK PG |
| D. linker | QRGASAGA IGG AKK RPK P |
| D.1 | KKRPKPGGGWNTGGSRYPGQG |
| D.2 | YPGQGSPGGNRYPPQGGGGWG |
| D.3 | GGGGWGQPHGGGWGQPHGGGW |
| D.4 | HGGGWGQPHGGGWGQPHGGG |
| D.5 | PHGGGGWGQGGTHSQWNKPS |
| D.6a | WNKPSKPKTNMKHVAGAAAA |
| D.6b | WNKPSKPKTNMKHMAGAAAA |
| D.7 | GAAAAGAVVGGLGGYMLGSA |
| D.8 | MLGSAMSRPLIHFGNDYEDR |
| D.9 | DYEDRYYRENMYRYPNQVYY |
| D.10 | NQVYYRPVDQYNNQNTFVHD |
| D.11 | NTFVHDCVNITVKQHTVTTTT |
| D.12 | VTTTTKGENFTETDIKMMER |
| D.13 | KMMERVVEQMCITQYQRESQ |
| D.14 | QRESQAYYQRGAS |

**Extended Data Table 2: Number of mice per group for the pooled urine**

| | 150 dpi | 250 dpi | 450dpi |
|---|---|---|---|
| Ddi | 7 mice<br>(All except #6) | 7 mice<br>(All except #6) | 3mice<br>(#2,3&4) |
| Mmo | 8 mice<br>(1 to 8) | 7 mice<br>(2 to 8) | 2mice<br>(#2 &4) |
| CPG | 7 mice<br>(1 to 7) | 7 mice<br>(1 to 7) | 1 mouse<br>#6 |

**Extended Data Table 3: Correlation of $PrP^C$ antibody titer and the percentage of positive replicates CWD seeding activity at different time point in feces**

| ID | 200 dpi | 250 dpi | 300dpi | 350dpi | End-point titer |
|---|---|---|---|---|---|
| Ddi 1 | 0 | 0 | ND* | ND* | 10,000 |
| Ddi 2 | 0 | 0 | 100 | 0 | 30,000 |
| Ddi 3 | 50 | 100 | 0 | 100 | ND# |
| Ddi 4 | 50 | 100 | ND | 50 | 30,000 |
| Ddi 5 | 0 | 0 | ND | 50 | 30,000 |
| Ddi 6 | 100 | ND | 100 | ND | 40,000 |
| Ddi 7 | 0 | 0 | 0 | 50 | 50,000 |
| Ddi 8 | 100 | 0 | 0 | 0 | 50,000 |
| Mmo 1 | 50 | ND* | ND* | ND* | 10,000 |
| Mmo 2 | 0 | 100 | 75 | 0 | 30,000 |
| Mmo 3 | 100 | 100 | 100 | 0 | 30,000 |
| Mmo 4 | 100 | 0 | 0 | 0 | 30,000 |
| Mmo 5 | 75 | 0 | ND | ND | 30,000 |
| Mmo 6 | 100 | 100 | 100 | 0 | 30,000 |
| Mmo 7 | 75 | 75 | 100 | 50 | 30,000 |
| Mmo 8 | 75 | 0 | 100 | ND | 5,000 |
| CPG 1 | 100 | 0 | ND | 100 | |
| CPG 2 | 50 | 50 | 100 | 50 | |
| CPG 3 | 50 | 100 | 100 | 100 | |
| CPG 4 | 100 | 100 | ND | 100 | |
| CPG 5 | 100 | 100 | 100 | 0 | |
| CPG 6 | 100 | 0 | 100 | 0 | |
| CPG 7 | 100 | ND | 100 | 75 | |
| CPG 8 | ND* | ND* | ND* | ND* | |

ND*: mouse euthanized before this time point, ND#: not enough sera collected and ND: feces can not be collected from this mouse. The scale ranges from 0 (all replicates were negative) to 100 (all replicates were positive).
